# Transcriptional dynamics of the oligodendrocyte lineage and its regulation by the brain erythropoietin system

**DOI:** 10.1101/2025.09.11.675672

**Authors:** Liu Ye, Vinicius Daguano Gastaldi, Yasmina Curto, Anne-Fleur Wildenburg, Xuan Yu, Martin Hindermann, Simone Eggert, Anja Ronnenberg, Qing Wang, Umer Javed Butt, Riki Kawaguchi, Daniel Geschwind, Wiebke Möbius, Susann Boretius, Manvendra Singh, Klaus-Armin Nave, Hannelore Ehrenreich

**Author notes:** Correspondence: Prof. Klaus-Armin Nave, PhD, Department of Neurogenetics, Max Planck Institute for Multidisciplinary Sciences, City Campus, Hermann-Rein-Str. 3, D-37075 Göttingen, Germany, Prof. Hannelore Ehrenreich, MD, DVM, Experimental Medicine, Department of Psychiatry and Psychotherapy Central Institute of Mental Health, Medical Faculty Mannheim, Heidelberg University, J 5, 68159 Mannheim, Germany. These authors contributed equally to this work (Liu Ye, Vinicius Daguano Gastaldi, Yasmina Curto, Anne-Fleur Wildenburg, Xuan Yu).

## Abstract

Oligodendrocytes differentiate from oligodendrocyte progenitor cells (OPC) in early postnatal development, but some oligodendrogenesis is maintained throughout adulthood, where oligodendrocyte lineage dynamics may contribute to neuroplasticity, adaptive myelination, and myelin repair. Here, we studied the effect of erythropoietin (EPO) and its receptor (EPOR) on oligodendrocyte lineage dynamics employing murine hippocampus and its myelinated fibers as model region. Using multiple stage-specific markers and single-nuclei-RNA-seq data, we find that EPO stimulates all oligodendroglial lineage cells directly, driving differentiation/maturation. Differential gene expression analysis reveals multiple EPO-regulated mRNAs, including downregulated transcripts for GABA-A receptors, fitting the known inhibition of oligodendrocyte maturation by GABA. Importantly, analogous oligodendrocyte responses are seen when endogenous EPO expression in brain is stimulated by hypoxia. Mice lacking EPOR from mature oligodendrocytes show subtle deficiencies of adult myelination in hippocampal fimbria and mild working memory deficits. These gain- and loss-of-function experiments may further suggest EPO as clinically safe treatment for remyelination therapies.

## INTRODUCTION

Oligodendrocytes (OL) wrap myelin for energy-efficient and rapid axonal impulse propagation. In addition, they provide metabolic support for axons and are thought to contribute to adult brain plasticity, adaptive myelination and cognitive performance^1, 2, 3, 4, 5, 6, 7^. Myelinating OL also contribute to cortical information processing^7^. The cognition-promoting properties of myelin resemble some of the effects of erythropoietin (EPO) on higher brain function and thus suggested a possible role of the brain EPO system in adult oligodendrogenesis and myelination. Next to its established physiological importance in neurons^8^, EPO might also directly influence oligodendrocyte lineage cells (OLC) in the adult brain. We therefore explored the direct responses of OLC to EPO and studied their functional significance.

EPO is a hypoxia-inducible growth factor, binding to a classical cytokine type 1 receptor, EPOR, and was named after its original description in erythropoiesis. However, EPO and EPOR are also expressed within the mammalian brain ^9, 10^. We previously discovered - by a ’reverse approach’ (human trials first) - that recombinant human (rh) EPO crosses the blood-brain-barrier and has potent procognitive effects. Importantly, these EPO effects in the brain are hematopoiesis-independent^11, 12, 13, 14^.

Searching for mechanistic insight in mice, we recognized that rhEPO markedly drives differentiation/maturation of pyramidal neurons from non-dividing progenitors in cornu ammonis (CA), a part of the hippocampus, outside known neurogenesis areas^15^. In this region, EPO strongly affects the transcriptome of basically all cell populations. Moreover, we noticed novel populations of newly differentiating pyramidal neurons, which are about 2-fold more abundant after rhEPO treatment with upregulation of genes crucial for neurodifferentiation, dendrite growth and spine density, synaptogenesis, memory formation, and higher cognition^8^. Subsequently, we detected that rhEPO restrains the inhibitory potential of interneurons in the murine hippocampus^16^, facilitating re-connectivity and synapse development. In parallel, we found that rhEPO dampens microglia activity and metabolism^17^.

We therefore hypothesized that also endogenous, i.e. brain-expressed, EPO acts in an auto/paracrine fashion and may have overlooked physiological significance, just imitated by rhEPO application. In fact, we demonstrated that the increased neuronal activity by motor-cognitive training leads to what we termed ’functional hypoxia’ which – similar to inspiratory hypoxia – stimulates endogenous EPO expression, particularly in neurons^17, 18, 19, 20, 21^. It appeared that EPO incites an integrated response of different brain cell types. In the neuronal circuitry, EPO thus has longer lasting effects, which we termed ’brain hardware upgrade’ ^21^. We now asked whether such an integrated response would also include OLC and adult myelination, and whether this would be caused by a direct OLC response to EPO.

We had previously observed that EPO administration to juvenile mice resulted, in addition to neuronal effects, in enhanced differentiation of preexisting OL progenitors, which was in the absence of DNA synthesis^15^. Moreover, in a behavioral screen with the IntelliCage paradigm^22^, *Cnp-Cre::Epor^flox/flox^* mice exhibited mild intellectual disability, but the underlying mechanisms remained completely unexplained. Other studies focused on rhEPO as a neuroprotective compound, reporting merely observations in the context of disease models, such as hypoxic-ischemic injury, stroke, and EAE ^23, 24, 25^. One recent paper investigated EPO/EPOR function in OL during early development, using conditional OPC-specific EPOR mutant mice^26^. However, the physiology and function of the EPO/EPOR system in the postnatal brain and in adult OLC has remained puzzling.

Therefore, the present work was undertaken to provide a comprehensive picture of adult EPO/EPOR functions in OLC of the cornu ammonis (CA1 and CA3). Research on fate specification, differentiation, and functional diversification of OLC primarily depends on the precise utilization of cell- or stage-specific molecular markers. We have used the conventional classification of OL progenitor cells (OPC, including early and late OPC), premyelinating OL (preOL or committed OL progenitors, COP), and the myelinating OL (MOL, comprising newly formed OL, NFOL, and mature myelinating OL, MOL1/2)^27^. To enhance the precision of our investigation into OL dynamics under EPO, we have undertaken a meticulous refinement of these stages through the incorporation of many different markers and their combinations. In addition, we employed our single nuclei transcriptome dataset of hippocampi from rhEPO versus placebo treated mice, in order to explore effects of rhEPO on the OLC transcriptional landscape. Our data lead us to discuss a potential involvement of the brain EPO system in adaptive myelination underlying higher cognitive performance.

## RESULTS

### Immunological markers and their combinations employed for OL classification

To investigate OLC dynamics under EPO, we conducted a detailed classification of known and recently described OLC stages, incorporating additional immunological markers and their combinations. An overview of all major stages together with their markers is presented in Figure 1a: (1) OPC, composed of early and late OPC, (2) preOL or COP, (3) MOL, comprising NFOL and MOL1/2. Platelet-derived growth factor receptor α (PDGFRα) and NG2 proteoglycan (also known as chondroitin sulfate proteoglycan4) are recognized OPC markers in the brain^28, 29, 30^. Since NG2 is also expressed in vascular pericytes, it is combined with the lineage-specific marker OL transcription factor 2 (OLIG2) to label OPC.

**Figure 1:**
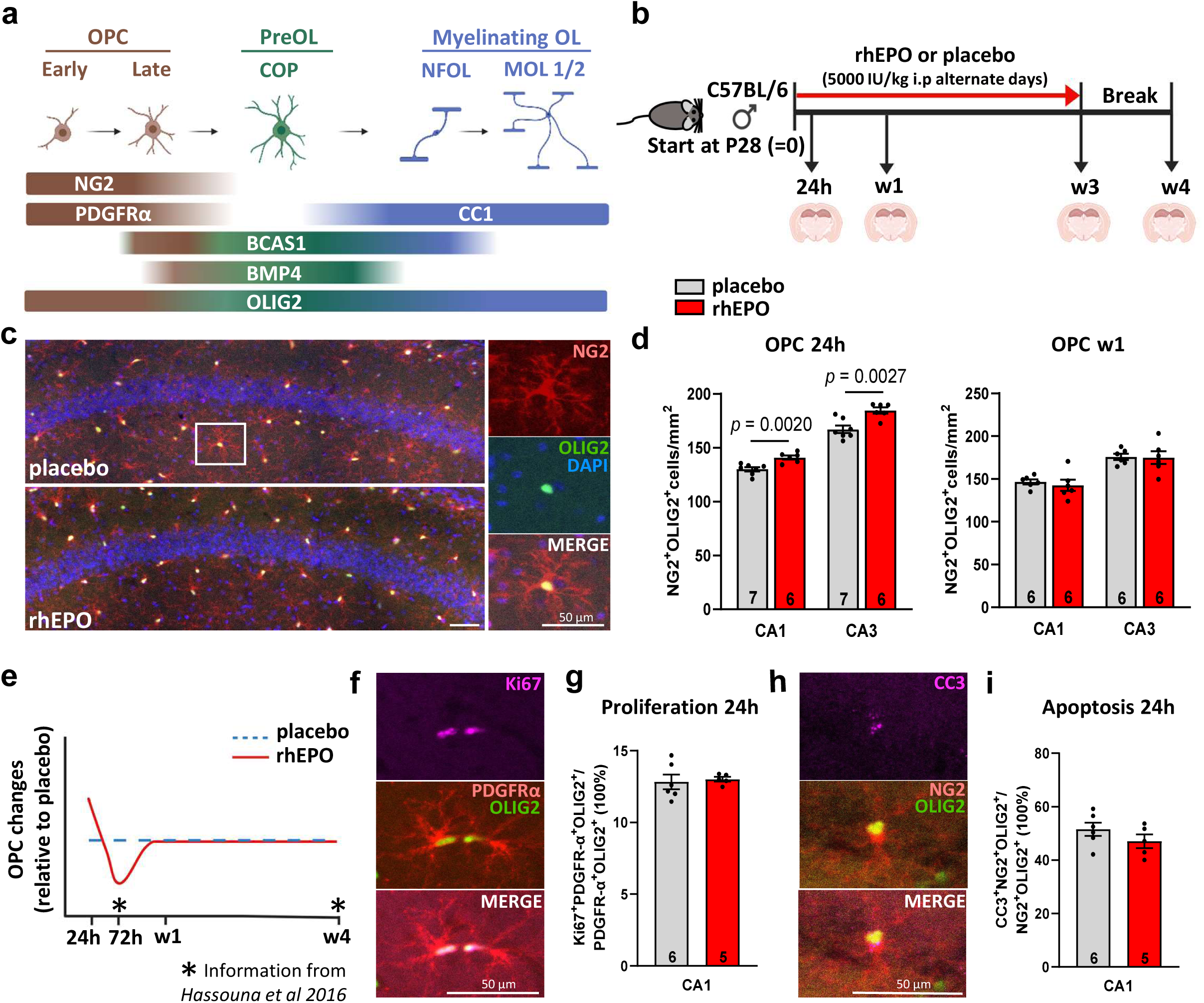
Effects of rhEPO/placebo on OPC numbers, proliferation, and apoptosis. **a** Schematic representation depicting OL differentiation and maturation, along with the respective stage markers. OPC, oligodendrocyte progenitor cells; PreOL, premyelinating oligodendrocytes; COP, committed oligodendrocyte progenitors; Myelinating OL, myelinating oligodendrocytes; NFOL, newly formed oligodendrocytes; MOL, mature myelinating oligodendrocytes. **b** Experimental design: Starting on P28, wildtype C57Bl/6 mice received rhEPO/placebo intraperitoneally, either once (24h experiment) or every other day during 1 week or 3 weeks. Mice were sacrificed 24h after the last injection, in the week 4 experiment after 1 week of break. **c** Representative images and **d** quantification of OPC (NG2^+^OLIG2^+^ cells) in CA1 and CA3 following 24h and 1 week of rhEPO/placebo, respectively. n = 7 mice (CA1 placebo), 6 mice (CA1 rhEPO), 7 mice (CA3 placebo), 6 mice (CA3 rhEPO) for the 24h timepoint; n = 6 mice for all groups at w1. **e** Sketch summarizing the course of OPC numbers at 24h, 72h, 1 week, and 4 weeks after initiation of rhEPO/placebo treatment. Information on the 72h and 4 week time point derived from Hassouna et al 2026. **f** Representative images and **g** percentage of proliferative OPC in CA1 at 24h. n = 6 mice (placebo), 5 mice (rhEPO). **h** Representative images and **i** % apoptotic OPC in CA1 at 24h. n = 6 mice (placebo), 5 mice (rhEPO). Scale bars in **c,f,h**: 50µm; mean±SEM presented; mouse n numbers given in bars; unpaired two-tailed Student’s t-test (**d,g,i**); **a, b, e** created with BioRender.com. Source data are provided as a Source Data file.

Engaging breast carcinoma amplified sequence1 (BCAS1) as further marker, we refine the OPC classification into early (NG2^+^BCAS1^-^OLIG2^+^) and late (NG2^+^BCAS1^+^OLIG2^+^) stages^31^. Notably, NG2^+^BCAS1^+^OLIG2^+^ cells develop distinct morphological characteristics with more intricate ramifications compared to NG2^+^BCAS1^-^OLIG2^+^ cells. PreOL or COP constitute terminally differentiated, immature OL, occupying an intermediate stage in OL progression between OPC and MOL/NFOL^32^. They have lost the prototypic OPC markers and initiate expression of major myelin genes, PLP (proteolipid protein 1), MBP (myelin basic protein), CNP (2′,3′-cyclic nucleotide 3′-phosphodiesterase, CNPase), MAG (myelin-associated glycoprotein), and MOG (myelin OL glycoprotein), primarily in cell bodies and proximal processes, while not yet forming compact myelin sheaths^33^. PreOL, with BMP4 and BCAS1 as markers, are at the root of OL differentiation and oligodendrogenesis. They present lower levels of cell cycle markers, while still expressing genes involved in migration *(TNS3, FYN)*^27, 34, 35^. These cells are highly branched and exhibit a morphologically complex profile, but reside at an immature state^36, 37^.

The MOL stage encompasses two distinct phases: NFOL can initiate the formation of myelin sheaths and gradually mature into fully developed OL. A key marker in this differentiation process is again BCAS1; its expression diminishes as OL progress into later stages of maturation, namely MOL1/2^31^. The monoclonal antibody anti-adenomatous polyposis coli (APC) clone CC1 is used to mark MOL without labeling myelin and allows counting mature OL. Combined with OLIG2, CC1 reliably labels MOL. Consequently, BCAS1^+^CC1^+^OLIG2^+^ cells are identified as NFOL, while BCAS1^-^CC1^+^OLIG2^+^ cells represent MOL1/2 population (Figure 1a).

### Influence of rhEPO on OPC

The rhEPO treatment scheme and timepoints of tissue harvesting are presented in Figure 1b. As early as 24h after a single rhEPO injection, OPC (NG2^+^OLIG2^+^) have increased in numbers in CA1 and CA3; this increase is temporary and has disappeared at week 1. A sketch illustrates the course of numerical OPC changes, with early expansion, followed by a decrease at 72h (as seen in ^15^) and a steady-state thereafter (Figure 1c,d,e). In fact, at the 72h time point, OPC undergo prominent differentiation. But what about the early enhancement? Under pathological conditions such as traumatic brain injury^38^ or experimental autoimmune encephalomyelitis in mice^25^, or in a rat model of embolic focal cerebral ischemia^39^, rhEPO has been found to increase OPC proliferation. During development (postnatal day 7), Muttathukunnel and colleagues^26^ observed also enhanced OPC proliferation in the subventricular zone of Tg21 mice. However, using Ki67 as proliferation and cleaved-caspase 3 (CC3) as apoptosis markers, we can exclude cell division or death upon rhEPO under physiological conditions in adolescent or young adult mice as explanation of the early increase *in vivo* (Figure 1f-i). In contrast, stimulating OPC in cell culture, we find at 24h augmented BrdU incorporation upon rhEPO in a dose-dependent fashion, peaking at 1IU/ml [with the bell-shaped curve showing a decrease after the peak (Figure 2a-c) as typical for many readouts after rhEPO application^40^]. This somewhat contradictory proliferation response *in vitro* is best explained by the essentially pure OPC cultures, where regulatory interactions with other cell types that may restrict proliferation *in vivo* are excluded^41^. There are several other possible explanations: (i) Differences in the proliferation markers used: In the *in vivo* experiments, we employed Ki67, a proliferative marker that labels cells in the G1, S, G2, and M phases, whereas in the *in vitro* experiments, we used BrdU, a proliferation marker, exclusively labelling S phase, hence providing perhaps a more precise snapshot. (ii) Delay in OPC proliferation *in vivo*: Although all experiments were conducted 24h after rhEPO administration, *in vitro* experiments are straightforward, with cells being directly stimulated by rhEPO and responding with both, proliferation and migration (Figure 2a-e), whereas *in vivo*, circulation and subsequent cellular effects may take more time.

**Figure 2:**
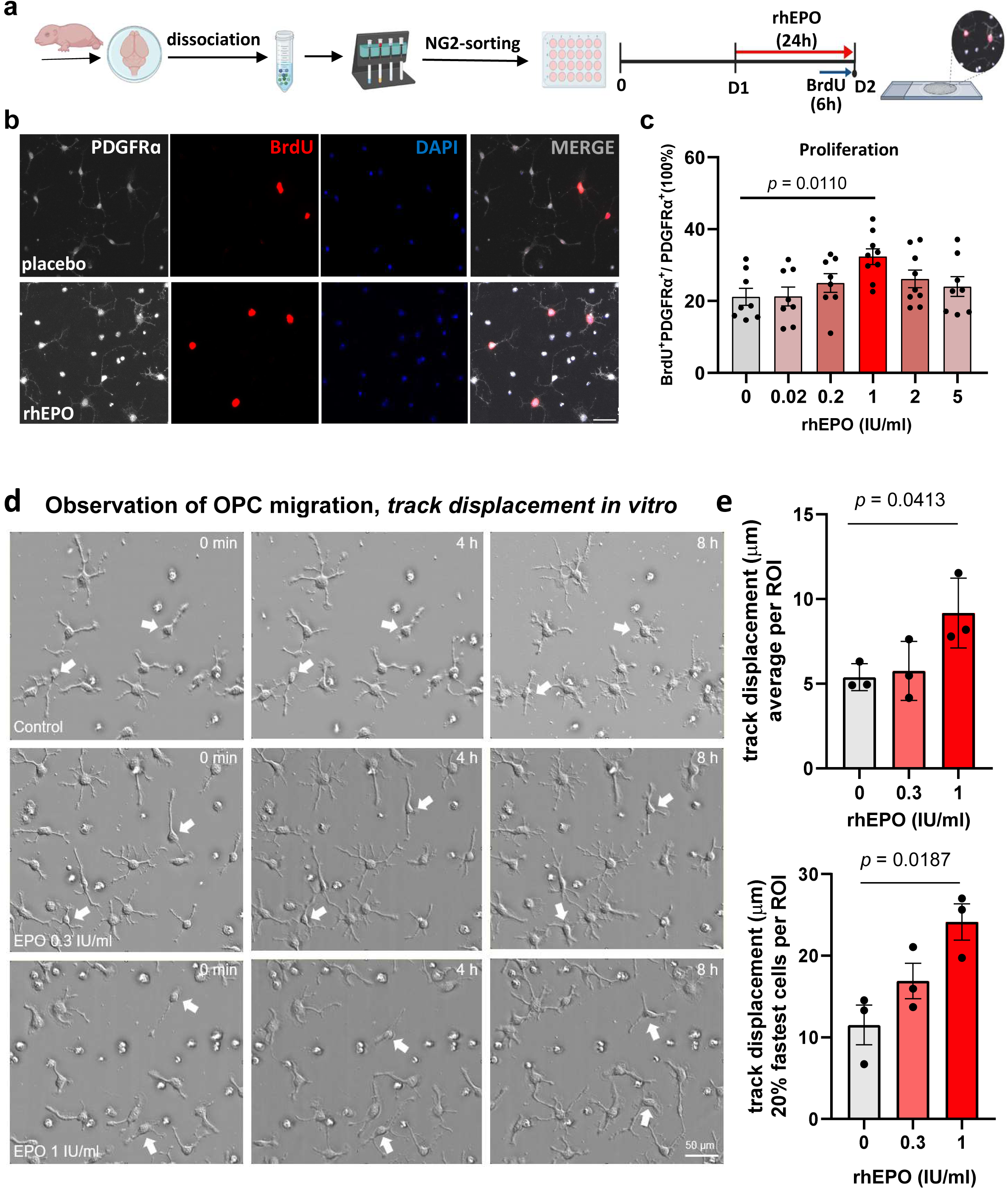
EPO induces OPC proliferation and migration *in vitro*. **a** Experimental design delineating the set-up of OPC cultures and subsequent *in vitro* procedures. **b** Representative culture images and **c** quantification of proliferating OPCs (BrdU⁺PDGFRα⁺ / PDGFRα⁺) after 24 h rhEPO treatment. Data were collected from three independent OPC cultures (biological replicates), each from pooled brains of P7 mice. For each condition, 8–9 fixed-area ROIs (≈2.5 mm²) were analyzed using FIJI. Each dot represents one ROI. **d** Primary OL were prepared via MACS sorting from C57/BL6N WT mice (P7) and imaged after 1 day of culture *in vitro* (DIV1) for 8h. Cells were treated with 0.3, 1 IU/ml rhEPO or an equivalent volume of the vehicle (EPREX). Representative images of the treated cells are shown. **e** Quantification of the migration distance of OPC given as “track displacement” for average values of all evaluated cells per ROI and underneath of the fastest 20% OPC per ROI. Data were collected from three independent OPC cultures (biological replicates). For each condition, 4 fixed-area ROIs (40x objective) were imaged and analyzed using Imaris. Scale bars in **b,d**: 50µm; mean±SEM presented; one-way ANOVA with Dunnett’s test (**c**), unpaired two-tailed Student’s t-test (**e**); **a**: created with BioRender.com. Source data are provided as a Source Data file.

### Effects of rhEPO treatment on the OL transcriptional landscape

#### Classification of clusters

Employing our snRNA-seq data sets, we followed the steps described in the Methods section. Our final data set for the analysis contained 12,043 nuclei that were unbiasedly classified into 5 clusters (Figure 3a; compare Figure 1a): OPC, COP, NFOL, MOL1/2. These clusters were defined using a combination of established OL marker genes (Figure 3b) and, to differentiate MOL1 and MOL2, pseudotime calculated using Monocle3^42^ (MOL2 pseudotime>MOL1 pseudotime; Wilcoxon Rank Sum test FDR adjusted p-value <0.001).

**Figure 3:**
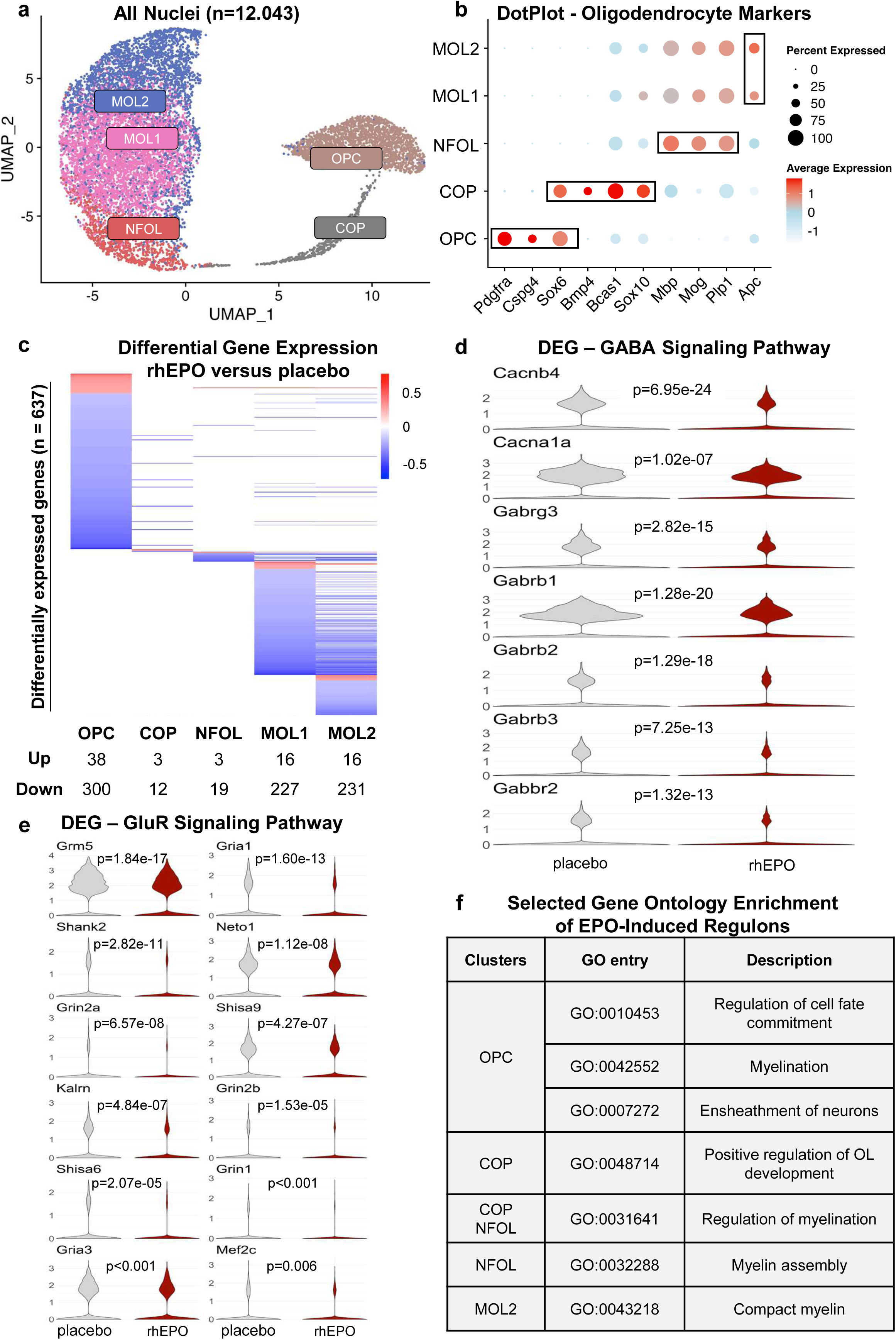
snRNA-seq analyses of OL upon rhEPO/placebo treatment. **a** Two-dimensional UMAP showing the 5 clusters obtained using unbiased transcriptome analysis of 12043 single nuclei, derived from hippocampi of mice treated with rhEPO/placebo. Each dot represents a single nucleus. Clusters were manually annotated by cross-referencing established markers (subfigure **b**). MOL1/2 were separated based on their calculated pseudotime (MOL2 pseudotime>MOL1 pseudotime, Wilcoxon rank sum test, p<0.001). **b** DotPlot illustrating the expression pattern of established OL markers. Average expression denotes the RNA expression of each marker normalized within respective clusters. These markers are used to confirm the identity of the clusters. MOL1 and MOL2 exhibit a higher level of *Apc* expression separating them from NFOL lineage. **c** Heatmap representing the differentially expressed genes (DEG, N=637) upon rhEPO/placebo for each cluster. The table beneath the plot shows the numbers of DEG for each cluster. DEG are represented by their average log2 fold change (minimum average log2FC ±0.25; adjusted p-value≤0.05, ‘bimod’ test, multiple correction done using Bonferroni). Red represents a higher expression value on rhEPO treated samples when compared to placebo samples, while blue represents a lower expression value for the same comparison. The full list of DEG is provided in Supplementary Table 1. **d** Multiple violin plots showing the expression level and distribution of DEG from the OPC cluster which belong to the Gene Ontology (GO) term *gamma-aminobutyric acid signaling pathway* (GO:0007214). Note that all DEG from this GO term were downregulated under rhEPO. P-values calculated using the ‘bimod’ test and adjusted using Bonferroni. Gray represents the expression values from placebo samples and red the ones from samples treated with rhEPO. **e** Multiple violin plots showing the expression level and distribution of DEG from the OPC cluster which belong to the GO term *glutamate receptor signaling pathway (*GO:0007215). Again, all DEG from this GO term were downregulated under rhEPO. P-values calculated using the ‘bimod’ test and adjusted using Bonferroni. Gray represents the expression values from Placebo samples and red the ones from samples treated with rhEPO. **f** Selected GO terms enriched significantly (over-representation analysis, adjusted p-value <0.05, Benjamini-Hochberg procedure, calculated using clusterProfile^57^) by the analysis of those genes that constructed the rhEPO exclusive regulons (see SCENIC analysis in Methods section). The full list of regulons and GO terms is provided in Supplementary Tables 3 and 4. Source data are provided as a Source Data file.

#### Differentially expressed genes (DEG)

Before starting our DEG examination, we removed one placebo sample and one EPO sample as previously described^8^. This was done to ensure the robustness of the analysis as these samples clustered together, separated of all others, when submitted to hierarchical clustering followed by 1000 bootstraps of relative gene expression levels^8^. The remaining samples underwent analysis as described in the Methods section. Briefly, nuclei were split according to treatment group, and compared across all 5 clusters using the ‘bimod’ test from Seurat^43^. A total of 637 unique DEG were identified, with the OPC cluster having the most, followed by MOL2 and MOL1 (Figure 3c). A complete list of all DEG is provided in Supplementary Table 1.

#### Gene ontology (GO) enrichment of the DEG

Next, we used WebGestalt^44^ to explore GO enrichment of the DEG in each cluster. Due to the low number of DEG in COP and NFOL, no GO terms were identified for these 2 clusters. Surprisingly, MOL2 also did not have any GO enrichment, while MOL1 showed only one (GO:0010498 - proteasomal protein catabolic process). On the other hand, the OPC cluster had a total of 64 enriched GO terms (complete list available in Supplementary Table 2). Here, we first highlight two specific GO terms, gamma-aminobutyric acid signaling pathway (GABA signaling pathway - GO:0007214, Figure 3d) and glutamate receptor signaling pathway (GluR signaling pathway - GO:0007215, Figure 3e). We note that changes of the GABA signaling pathway through its receptors have been connected to migration of OPC^45, 46^. Here, we found the reduction of 4 GABA-A receptor subunits (*Gabrg3*, *Gabrb1*, *Gabrb2*, *Gabrb3*). We also found the upregulation of matrix metalloproteinase *Mmp15* (average log2FC = 0.348, adjusted p-value = 0.005), a protein family previously connected to OPC migration^47^. We confirmed these findings *in vitro*, by measuring transcript levels of the GABA-A and glutamate pathways following addition of rhEPO to pure OPC cultures.

Somewhat unexpected GO terms for OPC include neuron projection organization (GO:0106027), axon development (GO:0061564), exocytosis (GO:0006887), and cognition (GO:0050890). Neuron projections include dendritic spines, which have their density increased after treatment with rhEPO^18^. The emergence of the axon development GO term in the OPC points to process outgrowth and the intricate relationship of these cells with neurons, as OPC receive synaptic input and have a role in fine-tuning of neural circuits^48^, but is mechanistically unexplained. Exocytosis may include the surface expression of signaling proteins. For OPC, this is of special interest also as exocytosis pathways play a role in oligodendrocyte development and myelination^49, 50^. And finally, cognition, specifically higher cognitive performance, is improved by rhEPO in multiple domains in both mice and humans^11, 12, 14, 51, 52, 53, 54, 55^. Whether these transcriptional changes also underlie the observed EPO effects awaits specific genetic analyses.

#### Gene regulatory networks (GRN)

Finally, we used SCENIC^56^ in an unsupervised manner to explore GRN between our treatments. Here we found only one regulon with *Anxa2* as a master regulator that was shared between OPC and COP. All other regulons were exclusive to a single cluster (available in Supplementary Table 3), suggesting that rhEPO acts differently on different OLC stages, including MOL1 and MOL2. After identifying these regulons, we used the genes that constructed them to calculate enrichment of GO terms using clusterProfiler^57^. A selection of GO terms from regulons that were exclusively induced by rhEPO is shown in Figure 3f. We note that rhEPO plays a role in differentiation and development early on, acting also in different stages towards ensheathment of neurons and myelination. These terms appear connected to multiple regulons (Supplementary Table 4), highlighting the broad and consistent action of rhEPO in myelination^15, 26, 58^. The complete list of the obtained GO terms is given in Supplementary Table 4.

### Impact of rhEPO on late OPC, PreOL/COP and NFOL/MOL1/2 in critical time windows

Next, we investigated dynamic changes of late OPC, PreOL/COP, and NFOL/MOL1/2 under rhEPO. Figure 4a-m shows that late OPC, PreOL/COP and NFOL respond to rhEPO with increased cell numbers in both CA1 and CA3, starting at 24h and extending to 1, 3 and 4 weeks. In fact, rhEPO affects OL differentiation in a “wave-like” pattern. At 24h (Figure 4b-d), an increase in late OPC/COP/NFOL (BCAS1^+^NG2^+^OLIG2^+^) is observed, while more mature BCAS1^+^NG2^-^OLIG2^+^ cells remain unchanged. By week 1 (Figure 4e,f), rhEPO leads to an increase in COP (BMP4^+^NG2^-^OLIG2^+^). At week 3 (Figure 4g-j), rhEPO primarily enhances the late OPC/COP population (BCAS1^+^CC1^-^OLIG2^+^). By week 4 (Figure 4k-m), rhEPO induces an increase in all late OPC/COP/NFOL populations. Finally, MOL reveal the upsurge not before week 4 (Figure 4n,o). Interestingly, the dentate gyrus displays an earlier increase of late OPC/COP/NFOL (i.e. at week 3, but no longer at week 4), suggesting non-cell-autonomous but region-specific differences *in vivo*. Indeed, MOL in the dentate gyrus did not show any increase in cell number at any time (Figure 5a-d).

**Figure 4:**
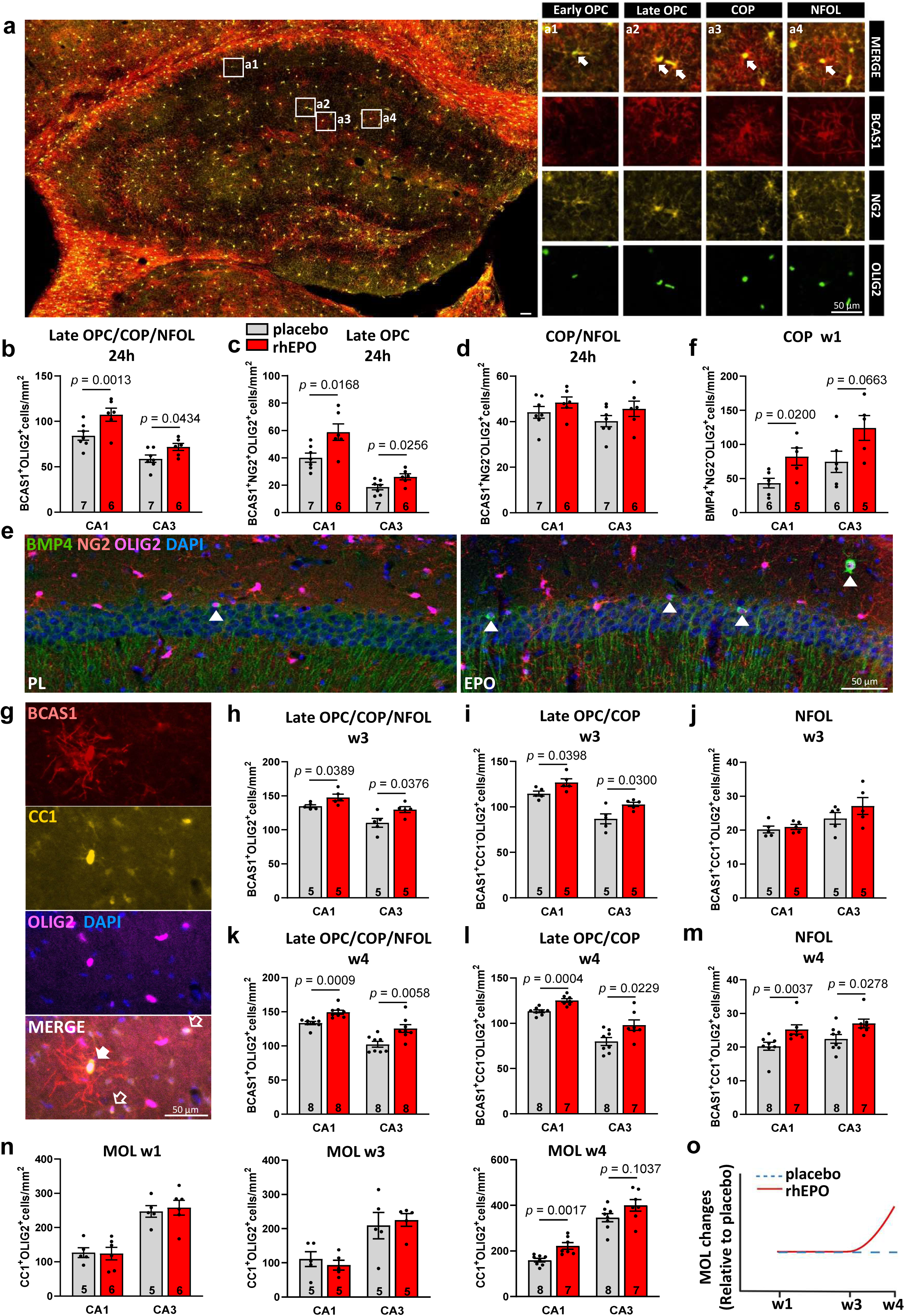
Effects of rhEPO on OL differentiation and maturation. **a** Overview of the hippocampus showing OPC/COP/NFOL marker BCAS1 (red), OPC marker NG2 (yellow), and OL marker OLIG2 (green) staining. Left: White frames denote areas **a1-a4**, magnified on the right. **a1** illustrates early OPC (BCAS1^-^NG2^+^OLIG2^+^), characterized by small soma with few ramifications; **a2** illustrates late OPC (BCAS1^+^NG2^+^OLIG2^+^) characterized by large soma with more ramifications; **a3** illustrates COP (BCAS1^+^NG2^-^OLIG2^+^) characterized by large soma with numerous ramifications; **a4** show NFOL (BCAS1^+^NG2^-^OLIG2^+^) characterized by large soma with parallel arranged ramifications; white solid arrows indicate early OPC, late OPC, COP and NFOL, respectively. **b-d** Quantification of late OPC/COP/NFOL (BCAS1^+^OLIG2^+^), late OPC (BCAS1^+^NG2^+^OLIG2^+^), and COP/NFOL (BCAS1^+^NG2^-^OLIG2^+^) in CA1 and CA3 after 24h of rhEPO/placebo. **e** Representative images displaying COP marker BMP4, OL marker OLIG2, and DAPI staining in CA1 after 1 week of rhEPO/placebo. White arrow heads indicate BMP4^+^NG2^-^OL. **f** Quantification of COP (BMP4^+^NG2^-^OLIG2^+^) in CA1 and CA3 after 1 week of rhEPO/placebo. **g** Representative images displaying late OPC/COP/NFOL marker BCAS1, MOL marker CC1, OL marker OLIG2, and DAPI staining. White solid arrow indicates NFOL (BCAS1^+^CC1^+^OLIG2^+^); white hollow arrows indicate MOL1/2 (BCAS1^-^CC1^+^OLIG2^+^). **h-j** Quantification of late OPC/COP/NFOL, late OPC/COP, and NFOL in CA1 and CA3 after 3 weeks of rhEPO/placebo. **k-m** Quantification in CA1 and CA3 of late OPC/COP/NFOL, late OPC/COP (BCAS1^+^CC1^-^OLIG2^+^), and NFOL (BCAS1^+^CC1^+^OLIG2^+^) at 4 weeks after initiation of rhEPO/placebo. **n** Quantification of MOL (CC1^+^ OLIG2^+^) at 1 week, 3 weeks and 4 weeks after initiation of rhEPO/placebo. **o** Sketch summarizing the course of MOL number changes at 1 week, 3 weeks and 4 weeks after initiation of rhEPO/placebo. Scale bars in **a,e,g:** 50µm; mean±SEM presented, n (number of mice) given inside the bars; unpaired two-tailed Student’s t-test, except for **b** (CA1), **m** (CA1), **n** (MOL W3, CA3), where two-tailed Mann-Whitney U test was performed. **o** created with BioRender.com. Source data are provided as a Source Data file.

**Figure 5:**
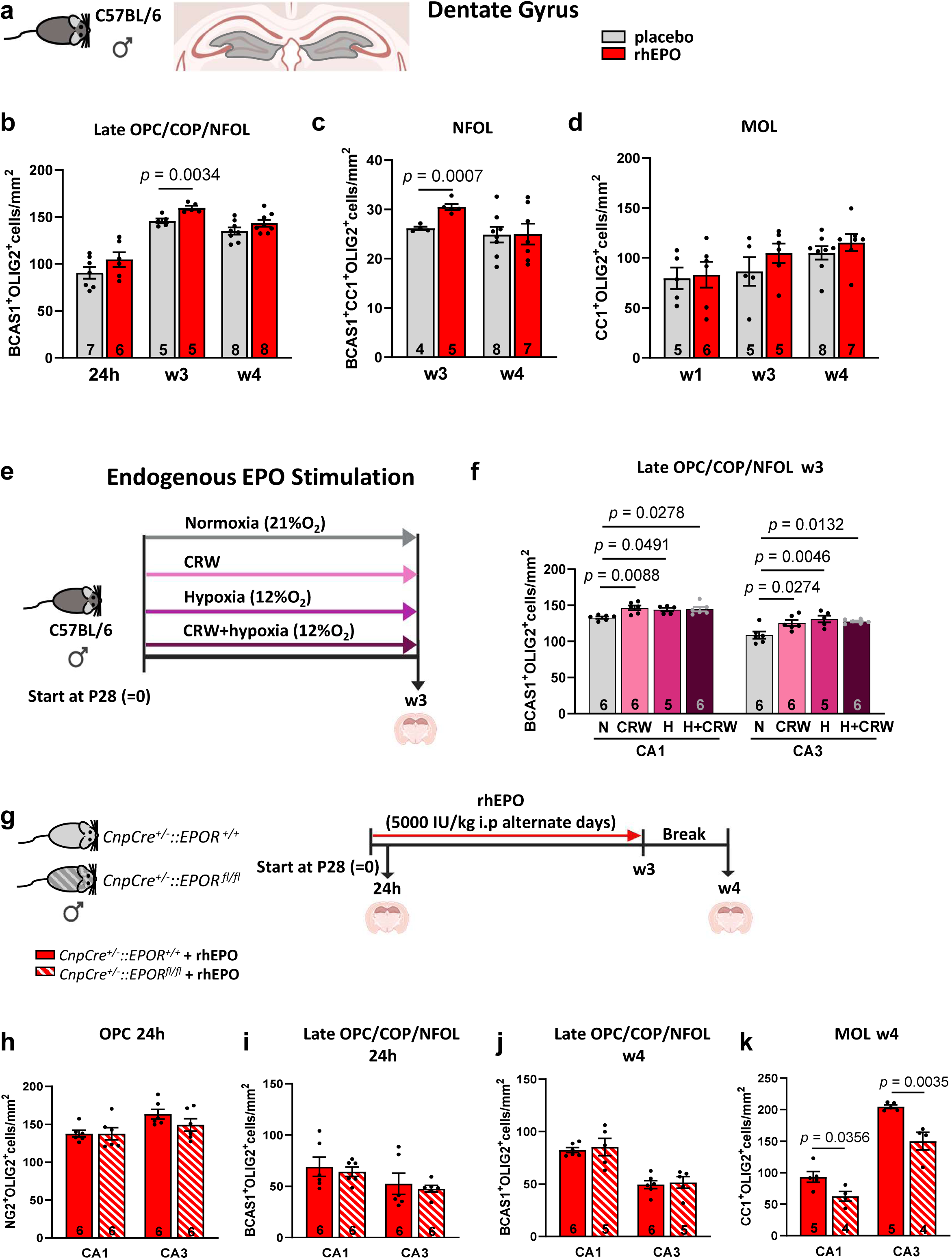
OL dynamics in dentate gyrus, induction of endogenous EPO, and consequence of EPOR deletion in MOL. **a-d Differential OL dynamics in the dentate gyrus** upon rhEPO/placebo. **a** Area of analysis. **b** Quantification of late OPC/COP/NFOL at 24h, 3 and 4 weeks following rhEPO/placebo initiation. **c** Quantification of NFOL at 3 and 4 weeks after start of rhEPO/placebo. **d** Quantification of MOL following 1 week, 3 weeks and 4 weeks after initiation of rhEPO/placebo. **e-f Induction of endogenous EPO** by motor cognitive challenge and inspiratory hypoxia. **e** Experimental design: Starting on P28, mice are exposed to normoxia (21% O_2_; N), functional hypoxia induced by complex running wheel (CRW) performance, inspiratory hypoxia (12% O_2_; H), or CRW upon inspiratory hypoxia (H+CRW), for 3 weeks. **f** Quantification of late OPC/COP/NFOL in CA1 and CA3. **g-k Effect of rhEPO in mice with EPOR deletion** in MOL. **g** Experimental design: *CnpCre::EPOR* mice received rhEPO once (24h) or for 3 weeks and brains were examined at week 4. **h** Quantification of OPC and **i** late OPC/COP/NFOL in CA1 and CA3 at 24h. **j** Quantification of late OPC/COP/NFOL and **k** MOL in CA1 and CA3 at 4 weeks; mean±SEM; n (number of mice) given inside the bars; unpaired two-tailed Student’s t-test except for **i** (CA3), where two-tailed Mann-Whitney U test was performed; in **f,** Tukey’s one-way ANOVA. **a,e,g** created with BioRender.com. Source data are provided as a Source Data file.

### Endogenous EPO, induced by ‘functional hypoxi’, increases late OPC, COP, NFOL

Since we observed strong effects of rhEPO on OL, we also tested the influence of endogenous, brain-expressed EPO, triggered by sustained neuronal activity (“functional hypoxia”), on OL dynamics. Employing male wildtype mice, we compared (i) functional hypoxia through motor-cognitive training with (ii) inspiratory (12% O_2_) hypoxia alone, and (iii) a combination thereof, starting on P28 and continuing over 3 weeks. All 3 treatments, known to stimulate brain EPO expression^17, 18, 19, 20, 21^ led to an increase in late OPC, COP, NFOL in CA1 and CA3 at week 3 (Figure 5e, f), similar to that observed after rhEPO (compare Figure 4h).

### Mice with EPOR deletion in MOL show subtle but distinct defects in adult myelin

To address the question whether rhEPO stimulates mature OL directly, we compared male *CnpCre^+/-^::EPOR^fl/fl^* mice, lacking EPOR in MOL, with *CnpCre^+/-^::EPOR^+/+^*mice as controls. Expectedly, rhEPO treatment of these mice resulted in analogous numbers of OPC, COP, NFOL in both groups at 24h and 4 weeks. Importantly, after 4 weeks, only CNP-expressing MOL (lacking EPOR expression) failed to respond to rhEPO (Figure 5g-k).

Next, we used a separate cohort of untreated *CnpCre^+/-^::EPOR* mutants and controls at the age of approximately one year. While female EPOR mutants underwent extensive behavior testing (see below) and had their brains studied immunohistochemically at age 50 weeks, male mutants at comparable age had magnetic resonance imaging (MRI) and afterwards electron microscopy (EM) completed (Figure 6a). Gross myelin quantification by MBP immunostaining in CA1 and CA3 did not reveal changes in EPOR deficient mice (Figure 6b,c). There were no signs of inflammation, i.e. microglia numbers were comparable between groups, as were activated microglia, OPC and MOL in CA1 and CA3 (Figure 6d-h). However, mice lacking EPOR in MOL present subtle defects of myelin in the hippocampal fimbria, with a shift in the size distribution of myelinated axons to the right. Thus, fimbria in the EPOR mutants have fewer myelinated small caliber axons, but g-ratios and the density of myelinated axons/µm^2^ are unaltered (Figure 6i,j). Although the Asymptotic Two-Sample Fisher-Pitman Permutation Test resulted in a significant p-value also for corpus callosum, we did not observe a systematic shift in the size of myelinated axons in the histogram (Figure 6k,l). Myelinated axons in CA1 are altogether rare and numbers were equivalent between groups (Figure 6m,n). However, MRI uncovered slight volume reductions in both fimbria and striatum of mutants (Figure 6o,p), whereas olfactory bulb, cortex, hippocampus, corpus callosum, and cerebellum do not differ from controls.

**Figure 6:**
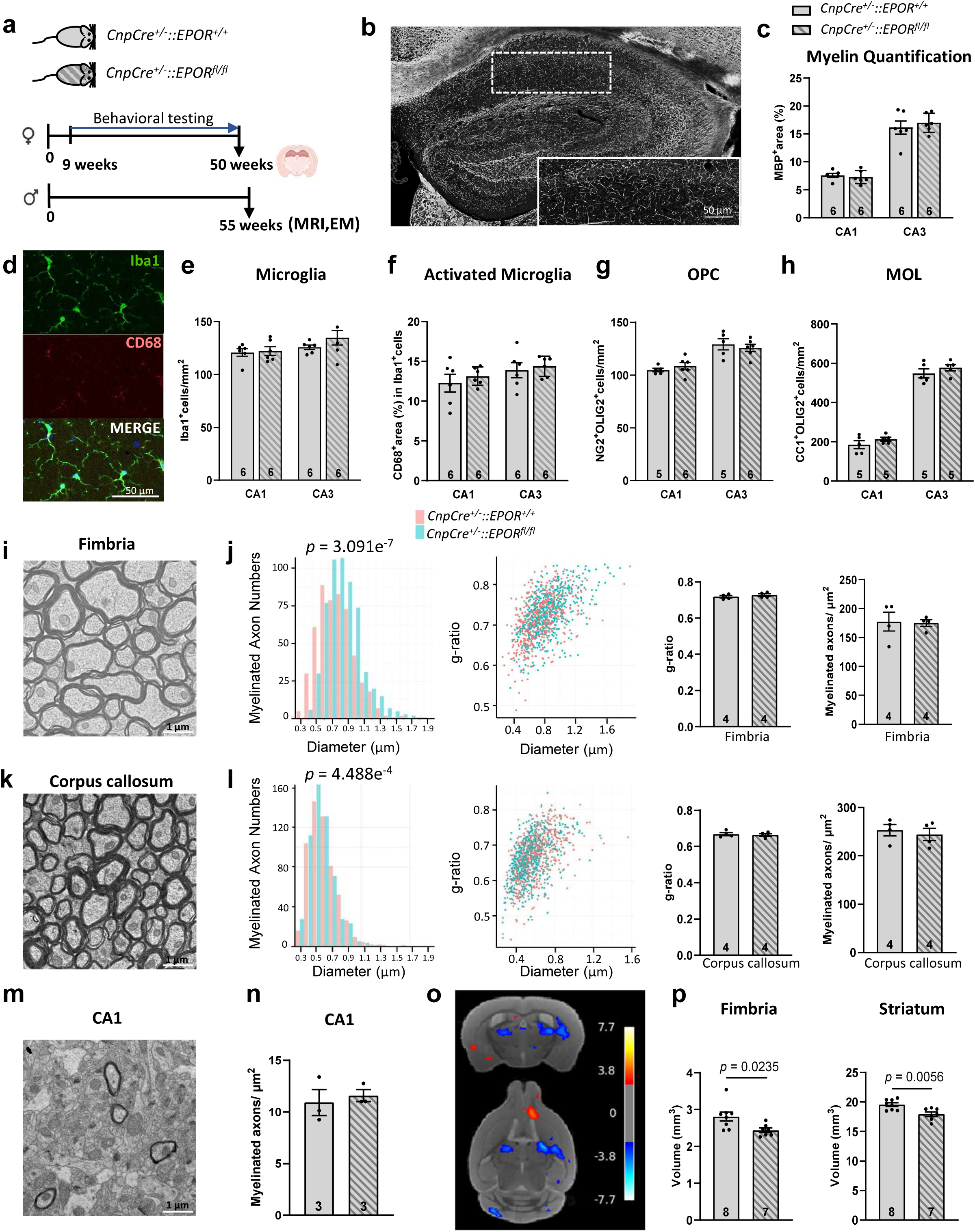
Deletion of EPOR in MOL has network-relevant influence on myelination. **a** Experimental design using *CnpCre::EPOR* mice and controls. **b** Representative images and **c** quantification of myelin (MBP^+^ area) in CA1 and CA3. **d** Representative images with microglia marker IBA1 and phagocytic (activation) marker CD68 and **e,f** quantification of microglia and activated microglia. **g,h** Quantification of OPC and MOL in CA1 and CA3 of *CnpCre^+/-^::EPOR^+/+^* and *CnpCre^+/-^::EPOR^fl/fl^* mice. **i** Representative EM images from fimbria of *CnpCre::EPOR* mice, showing no appreciable difference between genotypes. **j** Highly significant right-shift of myelinated axon calibers in fimbria, but no difference in myelin thickness (g-ratio) and myelinated axons/µm^2^. **k** Representative EM images from corpus callosum of *CnpCre::EPOR* mice, revealing no systematic alteration of myelinated axon calibers in fimbria, and no difference in myelin thickness (g-ratio) and myelinated axons/µm^2^. **m** Representative EM image from CA1 of *CnpCre::Epor* mice with **n** unchanged myelinated axons/µm^2^. **o** Comparison of Jacobian determinants in MRI of *CnpCre^+/-^::EPOR^fl/fl^* versus *CnpCre^+/-^::EPOR^+/+^* mice: red/yellow color indicates local tissue volume expansions, blue/cyan local reductions. **p** Volumetric MRI comparison of fimbria and striatum; scale bars: **b,d** 50µm, **I,k,m** 1µm; mean±SEM presented, with n (number of mice) given inside the bars; two-tailed Mann-Whitney U test was performed for **n**, asymptotic two-sample Fisher-Pitman permutation tests for **j** and **l**, unpaired two-tailed Student’s t-tests for the others. **a** created with BioRender.com. Source data are provided as a Source Data file.

### Mice with conditional EPOR deletion in MOL exhibit mild cognitive impairment

To assess the functional role of EPOR in MOL, female *CnpCre^+/-^::EPOR^fl/fl^* and control mice were subjected to an extensive behavioral battery comprising tests for anxiety, motor and sensory function, coordination, exploratory and social behavior, basic and higher cognition, including executive function and catatonic signs (Figure 7a). The main abnormalities were observed in higher cognitive performance, revealed with our sensitive IntelliCage design^59^. Unexpected alterations occurred in the anxiety profiles of *CnpCre^+/-^::EPOR^fl/fl^*mice, with abnormalities in elevated plus maze (increased time in open arms, p=0.049) and light-dark box (increased latency to enter the light compartment, p=0.03). Additionally, *CnpCre^+/-^::EPOR^fl/fl^*tended to an increased freezing behavior in basic (p=0.074) and pre-cue conditions (genotype: p=0.0133) of the fear conditioning test, suggesting a possible involvement of EPOR signaling and myelination in anxiety circuitries. For detailed overview of the behavioral phenotype see Supplementary Table 5.

**Figure 7:**
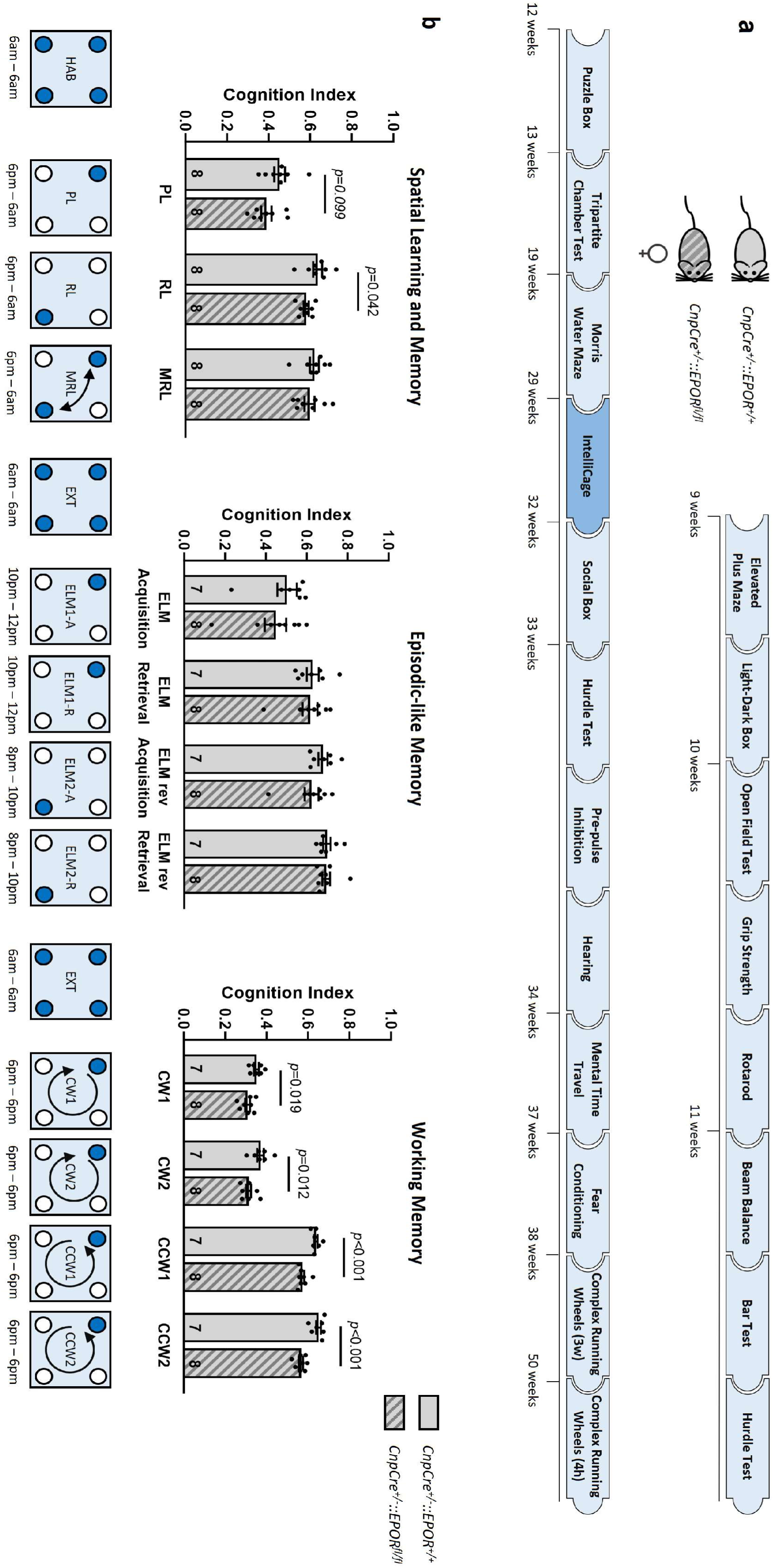
Mild cognitive impairment in mice with EPOR deletion in MOL. **a** Experimental outline of the behavioral test battery of *CnpCre::EPOR* mice. Behavioral characterization was started at the age of 9 weeks and completed at the age of 50 weeks. **b** Higher cognition was assessed in IntelliCage and cognition index per day in all challenges plotted. Below the graphs, a schematic outline of the IntelliCage paradigms is depicted; HAB: Habituation; PL: Place Learning; RL: Reversal Learning; MRL: Multiple Reversal Learning; EXT: Extinction; ELM: Episodic-like memory; CW: Clockwise; CCW: Counterclockwise; N numbers in bars; mean ± SEM; two-tailed two-sample t-test for normal and two-tailed Mann-Whitney U test for non-normal distributions. Only statistically significant p-values were shown, the non-significant values were 0.0993 (PL), 0.4795 (MRL), 0.2786 (ELM acquisition), 0.6007 (ELM retrieval), 0.2053 (ELM rev acquisition), 0.7789 (ELM rev retrieval). A more detailed overview of the results can be found in supplementary table 5. Source data are provided as a Source Data file.

Our IntelliCage paradigm, a comprehensive and standardized pipeline for automated profiling of higher cognition in mice^59^, consists of 3 challenges that assess different aspects of higher cognition. The cognition index was generated as a central readout of cognitive performance during these challenges. It does not only measure correct responses, but takes into account which errors a mouse would potentially make during a particular challenge and the relative weight of these errors^59^. Following habituation, challenge 1, extending over 3 nights, is the spatial learning and memory task. Mice have to learn which corner allows access to water (‘correct corner’), with the correct corner changing between nights (reversal learning) or changing every 3 hours (multiple reversal learning) (Figure 7b). *CnpCre^+/-^::EPOR^fl/fl^*mice showed a trend towards a reduced cognition index in the place learning task (p=0.099) and a decreased cognition index in reversal learning (p=0.042). Challenge 2 tests episodic-like memory. The challenge consists of 4 nights in which the mice have to identify the correct corner again, but this time during a specific timeframe, i.e. in night 1 from 10pm-12pm (acquisition). During night 2, this exact pattern is repeated (retrieval). Night 3 introduces a change of corner to the diagonally opposite corner, and a shift in the time slot to 8pm – 10pm (reversal – acquisition), which is repeated on night 4 (reversal – retrieval). No significant differences between groups were observed in this challenge. The last and most difficult challenge tests working memory and executive function, where the correct corner changes after every successful drinking attempt. On the first 2 nights, the corner changes clockwise, whereas on nights 3 and 4, the corners change counterclockwise. *CnpCre^+/-^::EPOR^fl/fl^* mice demonstrated reduced performance in this challenge with significantly lower cognition indices every night of the challenge (p=0.019, p=0.012, p<0.001, p<0.001, respectively). To conclude, mice lacking EPOR in MOL display dysfunctions in higher cognition, most likely as a consequence of subtle network disturbances.

## DISCUSSION

OL are best known for wrapping myelin, enabling energy-efficient and fast axonal impulse propagation. Moreover, OLC and adaptive myelination have been recognized as players in adult brain plasticity and cognitive performance ^1, 2, 3, 4, 5, 6, 7^. Neuroplasticity and cognition are also known targets of the neuronal EPO/EPOR system.

Here, we report that all cells of the oligodendrocyte lineage respond to exogenously applied rhEPO with differentiation and maturation towards MOL. These findings constitute an extension and refinement of previous knowledge. The lineage progression of rhEPO-responsive cells occurs in waves, with individual stage markers and marker combinations rising at subsequent time points. A similar pattern of pushed differentiation and maturation without DNA synthesis has been reported in excitatory neurons and their progenitors^8, 15^. Importantly, stimulation of endogenous EPO by functional or inspiratory hypoxia or their combination^21^ also leads to increases in OPC/COP/NFOL, similar to rhEPO treatment. This finding suggests a physiological role of the EPO system in OLC dynamics and we speculate that it might contribute to the widely studied phenomenon of adaptive myelination^60^ which is known to occur in response to neuronal activity.

Lack of EPOR in MOL leads to volume reduction in both fimbria and striatum, and to a right shift in the size distribution of myelinated axons of the hippocampal fimbria (when presented in the form of normal distributions), possibly explaining mild cognitive impairments. Future work will assess whether this mutation also affects adaptive cortical myelination in motor skill learning and other basic central nervous system (CNS) functions.

Of interest with respect to our clinical observations in EPO treated human patients^11, 12, 13, 14^ are the findings in *CnpCre^+/-^::EPOR^fl/fl^* mice regarding higher cognition. With our IntelliCage design^59^, such cognitive impairments can be readily identified, as demonstrated here, whereas conventional cognitive tests fail to show abnormalities. In line with our previous report (using a less sensitive IntelliCage paradigm), we observed a decreased cognition index in reversal learning^22^. Also in other dementia mouse models, impairments in reversal learning have been reported, with one study describing a more pronounced decline of reversal learning in female mice and suggesting a possible gender-specific vulnerability of cognitive flexibility^61, 62^. A particularly difficult challenge for EPOR mutants is the abrupt shift in the patrolling task from clockwise to counterclockwise. This task requires both working memory and executive functions which are known to depend on intact fornix and fimbria^63, 64^. It is possible that the mild dysmyelination of the hippocampal fimbria in EPOR MOL mutants is causally linked to the observed cognitive abnormalities. However, this assumption requires experimental validation.

We note that hippocampal dysfunction has been linked to anxiety-like behavior in mice, though with inconclusive results regarding their direction^65^. Altered anxiety profiles found in the current study, might reflect a perturbed hippocampal output.

Our single-nuclei-RNA-seq dataset^8^ revealed a plethora of transcriptional changes associated with oligodendroglial EPO/EPOR signaling, which makes it difficult to identify specific mechanisms and to prove causality. Exploring the candidate genes we found (see below) in more detail is close to impossible and we refrained from picking single genes, as their analysis would likely lead to premature take home messages.

However, we note mRNA changes that should be studied in simpler cellular systems in the future. This includes the upregulation of *Mmp15,* a matrix metalloproteinase^47^, and cell migration-associated genes, a potential effect of local EPOR signaling on (mitotic) OPC that are tiling the brain. We also found in OPC the reduction of 4 subunits of GABA-A receptors, which have been previously associated with OPC migration^45, 46^. Their downregulation may indicate a cessation of migration prior to differentiation, which needs to be confirmed in other systems. Also the enriched *Exocytosis* GO term in OPC^49^ has been linked to both migration and myelination^49,50^. Additionally, the downregulation of genes in the GO terms *GluR signaling pathway*, has been linked to OL differentiation before^45, 46^, and could link the effects of rhEPO to adult myelination^15, 26, 58^.

Other GO terms of interest in rhEPO stimulated OPC include *Exocytosis,* because direct communication with interneurons is essential for the maintenance of adequate social cognitive behavior^32^, a domain earlier shown to involve the EPO/EPOR system^66, 67, 68, 69, 70^. The GO term *Cognition* encompasses a broad number of genes/processes and here the role of myelination remains to be defined, but EPO/EPOR is well known to improve cognitive performance in rodents and humans^11, 12, 14, 51, 52, 53, 54, 55^. Finally, the GO term *proteasomal protein catabolic process* in MOL1 points to rhEPO stimulating myelin turnover and metabolic support function by MOL after myelin ensheathment ^71, 72^.

In conclusion, the brain EPO system and OL EPOR signaling profoundly influence OLC differentiation from early to late stages, OLC maturation and myelination. The OL EPOR contributes to adult neuroplasticity, complex network function and higher cognition, including working memory and executive performance. Given that rhEPO is a clinically approved and safe drug and that endogenous EPO can be stimulated by inspiratory as well as by functional hypoxia (motor-cognitive exercise), we assume that this knowledge will help in developing novel therapeutic strategies combatting cognitive disability and decline in the increasing number of neuropsychiatric conditions that are recognized to involve OL dysfunction.

## Methods

### Mice

All treatments and methods were conducted in accordance with the regulations of the local Animal Care and Use Committee (Niedersächsisches Landesamt für Verbraucherschutz und Lebensmittelsicherheit, LAVES) following the German Animal Protection Law (AZ 33. 19-42502-04-18/2803 & AZ 33.19-42502-04-17/2393 & AZ33.29-42502-04-22-00116). Every effort was made to minimize the number of animals used and their suffering. In all experiments, investigators were unaware of group assignment or treatment condition (‘fully blinded’).

All mice were group-housed in type IV cages (Techniplast Hohenpeiβenberg, Germany) inside ventilated cabinets (Scantainers, Scanbur Karlsunde, Denmark). They were segregated by both sex and genotype to prevent any potential aggressive interactions^22^. The cages were furnished with wood-chip bedding and nesting material (Sizzle Nest, Datesand, Bredbury, United Kingdom). All mice were housed in a temperature-controlled environment (21±2°C, humidity ∼50%) on a 12h light/dark cycle. They had access to food (Sniff Spezialdiäten, Bad Soderberg, Germany) and water *ad libitum,* except for the female *CnpCre::EPOR* mice during IntelliCage testing (see below).

Different cohorts of juvenile C57BL/6N male mice were used at the starting age of P28 for the different rhEPO treatment studies described below. Additionally, *CnpCre::EPOR* mice were generated to specifically delete *Epor* in mature OL. Briefly, *EPOR^fl/fl^*mice^18^ were bred with *CnpCre* mice^73, 74^ to generate *CnpCre^+/-^::EPOR^fl/fl^* (OL EPOR KO) and their wildtype littermates (*CnpCre^+/-^EPOR^+/+^*). Young *CnpCre::EPOR* male mice of both genotypes were used for immunohistochemical (IHC) analysis after treatment with rhEPO starting at P28. An additional cohort of untreated *CnpCre::EPOR* female mice were employed for behavioral studies, followed also by IHC. Untreated *CnpCre::EPOR* male mice of matching age (55 weeks) were used for MRI and afterwards electron microscopy (EM).

### Treatments

#### Recombinant human (rh) EPO

(5000 IU/kg body weight; NeoRecormon, Roche, Welwyn Garden City, UK) or placebo (solvent control solution, EPREX® buffer) were applied via i.p. injections (0.01ml/g) either once or every other day as indicated in the treatment schemes of figures, always starting on P28. Different sets of C57BL/6N male mice followed rhEPO/placebo treatment either once (24h experiment) or every other day during 1 week (w1 experiment) or 3 weeks (w3 experiment). All mice were sacrificed 24h after the last injection, in the w4 experiment at 1 week after 3-week rhEPO/placebo treatment. C57BL/6N mice used for snRNA-seq study underwent the same 3-week rhEPO/placebo treatment starting at P28. *CnpCre::EPOR* mice were treated with rhEPO and sacrificed after 24h of a single rhEPO injection (24h experiment) or one week after cessation of the 3-weeks rhEPO treatment (week 4 experiment).

#### Stimulation of endogenous EPO expression

For **inspiratory hypoxia experiments**, mice were put into a large, custom-designed hypoxia chamber (Coy Laboratory Products, Grass Lake, MI, USA) at 12% O_2_ for 3 weeks from P28 to P49, starting with a gradual reduction of 3% O_2_ per day for 3 days. **Complex running wheels** [CRW^18^; TSE Systems, Bad Homburg, Germany] for voluntary running were placed into cages for 3 weeks from P28 to P49. Running was voluntary at all times with *ad libitum* access to food and water. Groups included (1) normoxic room conditions (at 21% O_2_) in standard cages (N group), (2) normoxic conditions with voluntary running on CRW (CRW group), (3) hypoxic conditions (hypoxia chamber to 12% O_2_) in standard cages (H group), and (4) hypoxic conditions with voluntary running on CRW (H+CRW group). Mice were sacrificed, perfused, and brains collected as described below.

#### Immunohistochemistry (IHC)

Except otherwise indicated, mice were anesthetized by i.p. injection of Avertin and transcardially perfused via left cardiac ventricle with Ringer’s solution followed by 4% paraformaldehyde (PFA) in sodium phosphate-buffered saline (PBS) 0.1 M, pH 7.4. Dissected brains were post-fixed in 4% formaldehyde and equilibrated subsequently in 30% sucrose dissolved in PBS at 4°C overnight. Brains were then embedded in cryoprotectant (O.C.T.TM Tissue-Tek, Sakura) and stored at −80 °C. Whole mouse brains were cut into 30μm thick coronal sections with a Leica CM1950 cryostat (Leica Microsystems, Wetzlar, Germany) and stored at -20°C in 25% ethylene glycol and 25% glycerol in PBS until use. Following blocking and permeabilization with 5% normal horse serum (NHS) in 0.3% Triton X-100 in PBS (PBST) for 1h at room temperature (RT), primary antibodies were incubated in 5% NHS with 0.3% PBST over 2-3 nights at 4°C. The following primary antibodies were used in this study: anti-AN2/NG2 (rat, 1:1000,^75^), anti-OLIG2 (rabbit, 1:1000, AB9610; Millipore-Sigma; goat, 1:1000, AF2418; R&D Systems), anti-PDGFRα (rabbit, 1:1000, #3174; Cell Signaling Technology), anti-Ki67 (rat, 1:500, 14-5698-82; eBioscience), anti-cleaved caspase3 (rabbit, 1:200, #9664; Cell Signaling Technology), anti-BACS1 (rabbit, 1:1000, 445003; Synaptic Systems), anti-BMP4 (rabbit, 1:100, ab39973; Abcam), anti-APC/CC1 (mouse, 1:500, OP80; Millipore-Sigma), anti-MBP (rabbit, 1:1000, custom-made in the Nave Laboratory), anti-IBA1 (chicken, 1:1000, 234006; Synaptic Systems), anti-CD68 (mouse, 1:500, FA-11; Bio-Rad). After washing, sections were incubated for 2h at room temperature with different secondary antibody cocktails diluted in 3% NHS with 0.3% PBST. The following fluorescently conjugated secondary antibodies were used: Donkey anti-goat Alexa Fluor-488 (1:500, A11055; Thermo-Fisher), donkey anti-rabbit Alexa Fluor-488 (1:500, A21206; Thermo-Fisher), donkey anti-chicken Alexa Fluor-488 (1:1000, 703–545-155; Jackson ImmunoResearch), donkey anti-rat Alexa Fluor-555 (1:250, ab150154; Abcam), donkey anti-mouse Alexa Fluor-555 (1:500, A31570; Thermo-Fisher), donkey anti-goat Alexa Fluor-555 (1:500, A21432; Thermo-Fisher), goat anti-rat Alexa Fluor-555 (1:1000, A21434; Thermo-Fisher), donkey anti-rabbit Alexa Fluor-647 (1:500, A31573; Thermo-Fisher), goat anti-rat Alexa Fluor-647 (1:500, A21247; Thermo-Fisher). Nuclei were stained for 10min at RT with 4′,6-diamidino-2-phenylindole (DAPI; 1:5000; Millipore-Sigma) in PBS. Finally, sections were washed in PBS 0.1 M and mounted on SuperFrostPlus Slides (ThermoFisher) with Aqua-Poly/Mount (Polysciences, Inc).

#### OL Progenitor Cell (OPC) Culture

Magnetic-activated cell sorting was performed to isolate NG2-positive OPC (with anti-AN2 MicroBeads, Miltenyi Biotec, 130-097-170) from P7 wildtype C57BL/6N mouse brains. Brain tissues were quickly isolated and cut into small pieces on ice after removing blood vessels. Mouse brain dissociation was performed using the neural tissue dissociation kit (Miltenyi Biotec,130-092-628) following manufacturer’s protocol. The GentleMACS™ octo dissociators with heaters (Miltenyi Biotec, 130-096-427) were applied for the mechanical dissociation steps. After the dissociation steps, cell suspension was centrifuged at 238 × g for 10min. The pellet was resuspended and incubated in warm OPC culture medium (100ml NeuroMACS media, 2ml MACS NeuroBrew21, 1ml Penicillin/Streptomycin, and 1ml L-GlutaMAX), rotating for 2h at 37°C. Cell suspension was cooled down by centrifuging at 238 × g, 4°C for 10min. The pellet was re-suspended and incubated with pre-cold DMEM with 1% horse serum and mixed with anti-AN2 microBeads (10μl AN2-beads per 10^7^ total cells) for 15min at 4°C. Cell suspension flowed through the pre-activated LS columns (Miltenyi Biotec) which were attached on the QuadroMACS™ Separator (Miltenyi Biotec). LS columns were detached after being washed 3 times with 3ml DMEM containing 1% horse serum. Cells that combined with AN2 beads were collected by flushing the columns with 5ml DMEM containing 1% horse serum. The cell suspension was centrifuged and re-suspended with OPC culture medium with 4.2µg/ml Forskolin, 0.01µg/ml CNTF, 0.01µg/ml PDGF and 0.001µg/ml NT3. OPC were plated at a density of 6 x10^4^ cells per well on a 24-well plate with coverslips in a proliferation medium. After 24h, cells were treated with 0.02, 0.2, 1, 2, 5IU/ml rhEPO or an equivalent volume of solvent solution (placebo). After 18h, BrdU (Abcam, ab142567) was added to the cells (1µl 20mM BrdU per well), which were incubated for another 6h.

#### Live Cell imaging

OPC were prepared as described above from P7 wild-type C57BL/6N mouse brains and seeded at a density of 50.000 cells per well in proliferation medium in an 8 -well glass bottom live cell chamber (Ibidi 80827). After 24h, cells were treated with 0.3IU/ml or 1IU/ml rhEPO or an equivalent volume of the vehicle (EPREX® buffer) and imaged for 8h. Differential interference contrast (DIC) images were acquired with an inverse Nikon Eclipse Ti2 microscope (Nikon, Tokyo, Japan) equipped with 40x (0.95 NA) air objective and a Prime95B camera (Teledyne Photometrics). The Ibidi dish was held at 37°C, 5% CO_2_ during imaging with live cell imaging components from Okolab (www.oko-lab.com). The software used for imaging was NIS-Elements 4.5. Cell movement of the OPC culture was evaluated using the cell tracking tool of the Imaris software 10.1.1 (Bitplane). The cells were analyzed using ‘autoregressive motion’. ‘Track Displacement Length’ was selected for quantification. The tracks generated were checked in the ‘Surpass Viewer’ and the resulting data were statistically analyzed via Prism 10.3.1.

#### Immunocytochemistry (ICC)

Cells were fixed with 4% PFA and washed with PBS 3x for 5min each, followed by permeabilization with cold PBS containing 0.3% Triton X-100 and blocked with 10% goat serum and 0.03% Triton X-100 in PBS for 1h. The primary antibodies anti-BrdU (rat, 1:1000, ab6326; Abcam), and anti-PDGFRα (rabbit, 1:1000, ab203491; Abcam) were diluted in 1.5% horse serum in PBS and applied at 4°C overnight. Secondary antibodies donkey anti-rat Alexa Fluor-555 (1:1000, A48270; Thermo-Fisher); donkey anti-rabbit Dylight650 (1:1000, SA5-10041; Thermo-Fisher) were diluted in PBS and applied at room temperature for 1h after washing the cells 3x with PBS. Coverslips were washed again and exposed to DAPI in PBS for 10min. Cells were mounted with Aqua-PolyMount, waiting for imaging.

#### Microscope Imaging and Analyses

For analysis of NG2, OLIG2, PDGFRα, Ki67, cleaved caspase3 (CC3), BCAS1, and CC1, a Nikon Ti2 Eclipse (Nikon, Tokyo, Japan) epifluorescent microscope with 40x objective (NA0.6) was used. For BMP4, NG2, OLIG2, MBP, IBA1, and CD68 quantification, sections of hippocampus were acquired as tile scans on a confocal laser scanning microscope (LSM 880, Zeiss), equipped with a 20x air objective (20×/0.8 M27). Z-Stack scans were performed at 3 µm intervals for BMP4, NG2, and OLIG2 staining. Quantifications and image processing were performed with FIJI-ImageJ software^76^. NG2^+^OLIG2^+^ cells (OPC), Ki67^+^PDGFRα^+^OLIG2^+^ cells (proliferating OPC), CC3^+^NG2^+^OLIG2^+^ cells (apoptotic OPC), BCAS1^+^OLIG2^+^ cells (late OPC/COP/NFOL), NG2^+^BCAS1^+^OLIG2^+^ cells (late OPC), BMP4^+^NG2^-^OLIG2^+^ cells (COP), CC1^+^OLIG2^+^ cells (MOL), BCAS1^+^CC1^+^OLIG2^+^ cells (NFOL), and IBA1^+^ cells (microglia) were manually counted. MBP^+^ area and CD68^+^ area in IBA1^+^ cells were quantified densitometrically upon uniform thresholding. Cell counts, MBP^+^ area, and CD68^+^ area in IBA1^+^ cells were normalized to quantified areas (CA1, CA3, dentate gyrus). Data from 3 to 5 hippocampal sections/mouse was averaged.

For the OPC culture experiment, cells were imaged after PDGFRα and BrdU staining by a bright-field light microscope (Zeiss AxioImager Z1, coupled to Zeiss AxioCam MRc Camera) at 20x magnification, controlled and stitched by ZEN imaging software (ZEN Blue v3.3, Zeiss). Biological replicates (3 from independent cultures) were performed and 4 non-overlapping random images per group were chosen from each replicate. PDGFRα^+^BrdU^+^ cells/PDGFRα^+^ cells were counted.

#### Transponder Placement

The transponder placement surgery was performed as described before^59^. In short, female *CnpCre::EPOR* mice were anesthetized by intraperitoneal injections of 2,2,2-tribromoethanol (1.36%, T48402, Sigma) in ddH_2_O (Avertin, 0.3 mg/gBW). In the skin of the neck, a small incision of ∼2mm was made to allow implanting of ISO standard transponders (length: 8.5mm; diameter: 1.2mm; PM162-8, PeddyMark, Redhill, United Kingdom). Sutures closed the incision, and mice were put in their home cage on a heating plate (∼38°C). The transponders allow for experimenter-independent tracking in the IntelliCage paradigms (TSE Systems, Bad Homburg, Germany).

#### Behavior Battery

At the age of 9 weeks, female *CnpCre::EPOR* mice started a behavioral testing battery, including tests for domains such as exploratory and anxiety-related behavior, motor and sensory functions, higher cognition, and social conduct. The behavioral tests were performed in the following order: Elevated Plus Maze, Light-Dark Box, Open Field Test, Grip Strength, Rotarod, Beam Balance, Bar Test, Hurdle Test, Puzzle Box, Tripartite Chamber Test, Morris Water Maze, IntelliCage, Pheromone-Based Social Box, Hurdle Test, Pre-Pulse Inhibition, Hearing Test, Mental Time Travel, Fear Conditioning, and Complex Running Wheels. Upon completion of the behavioral battery, mice were perfused and brains utilized for IHC. Unless otherwise indicated, experiments were performed in the light phase. During the course of the experiments, animals that were unable to perform were excluded from the test.

#### Elevated Plus Maze

Elevated Plus Maze was performed as previously described^77^. In short, mice were placed in the center of a gray Perspex plus-shaped arena (light intensity: 129 lux), which comprises a central platform (5x5 cm), and two open and two walled arms (each 30x5cm, and 30x5x15cm respectively). Mice were allowed to freely explore the arena for 5 minutes, while automated tracking software (Viewer3, Biobserve, Bonn, Germany) recorded the total distance [cm], as well as time spent [s] per zone (open vs. walled arms). Animals were tested at the age of 9 weeks.

#### Light-Dark Box

The light-dark box (35x20x19cm) consists of an open, brightly illuminated compartment (light intensity: 124 lux), and a closed, dark compartment, divided by a black partition. An opening in the partition allows the mice to travel between compartments. Mice were placed in the dark compartment, and allowed to explore the arena for 5 minutes. A cut-off value of 180 seconds was used for mice not entering the light compartment within that time. Latency to enter the light compartment [s] was manually scored. Animals were tested at the age of 9 weeks.

#### Open Field Test

The Open Field Test was performed as previously described^77^. Briefly, mice were placed in the center of a gray Perspex arena (120 cm diameter, 25 cm height of wall, light intensity: 136 lux), after which time to reach the outer wall [s] was manually assessed by the experimenter. If mice did not reach the outer wall within 180 seconds, the experimenter manually placed the mouse in the peripheral zone. Afterwards, mice were allowed to freely explore the arena for 7 minutes. Automated tracking software (Viewer3, Biobserve, Bonn, Germany) recorded the total distance [cm], as well as the time spent per zone (center, intermediate, and peripheral) [s]. Animals were tested at the age of 10 weeks.

#### Grip Strength

To assess forelimb strength, a grip strength meter as described previously^77^ was used, which consists of a metal bar connected to a measurement device. Mice were lifted by their tails and brought near the rod, to allow their forepaws to grasp the rod. When the rod was grasped, the mice were gently pulled backward by their tail until the rod was released. The maximum of exerted grip strength [au] was recorded by the device. Per animal, three trials were done, with an average of those three trials used for further statistical analysis. Animals were tested at the age of 10 weeks.

#### Rotarod

The Rotarod test was performed as described in detail by Netrakanti et al (2015)^77^. In short, animals were placed on a rotating drum, that accelerates from 4-40 rounds per minute (RPM) over the course of 5 minutes. This test was preceded by a habituation time to the non-moving rod. The latency to fall off [s] is recorded for each mouse. To assess motor learning, a second test was performed 24 hours later. Animals were tested at the age of 10 weeks.

#### Beam Balance

Netrakanti and collaborators (2015) provided a detailed description of the Beam Balance apparatus and test^77^. In short, mice were placed on a horizontal beam with an illuminated start (light intensity: 125 lux) and a dark compartment to seek shelter at the other end of the beam. The test takes three days, with two phases each. Beam thickness is reduced during the course of the phases. On the first day, mice were first placed on the 25mm beam, directly in front of the dark cage. During the second trial, they were placed in the middle of the 25mm beam. In the following trials, mice were placed at the illuminated start. The second day comprised a habituation trial on the 25mm beam, after which a trial with a reduced beam thickness of 10mm was performed. The third day followed a similar design with a habituation to 25mm and a second trial on an 8mm beam. The escape latency [s] was recorded for both trials on the second day and the trial with the 8mm beam on the third day. If mice do not succeed on the first trial of a phase, the phase will be repeated to a maximum of three trials.

#### Bar Test

The bar test was used to assess the presence of catatonic signs^78^. In this test, a mouse is gently raised by its tail and brought near a horizontal steel bar. The mouse was allowed to grip the bar with its forepaws while the hind paws touched the ground, after which the tail was released. Each mouse underwent two consecutive trials. A trial is considered invalid if the mouse fails to have both forepaws at the bar and both hind paws on the floor simultaneously. The tests were recorded using a high-resolution camera, and time immobile at the bar [s] was manually scored. Two independent raters scored the recorded videos. The scoring was applied to the first trial, and in case of invalidity, the second trial was scored. If both trials were deemed invalid, the mouse was excluded from further analysis for this test. Animals are tested at the age of 11 weeks.

#### Hurdle test

To test for executive function in catatonia, the hurdle test was developed^79^. The mouse was placed in the center of a circular arena (120 cm diameter, 25 cm height of wall, light intensity: 140 lux), in which a PVC comb insert consisting of interconnecting hurdles (10x10 cm) was placed. Driven by their motivation to avoid brightly illuminated and open spaces, mice strive to reach the periphery as fast as possible, after which the test was stopped. Each mouse underwent two trials. To assess motor learning, the test was repeated six months later. Automated tracking software (Viewer3, Biobserve, Bonn, Germany) recorded the latency to reach the periphery [s]. Latency to reach the periphery and number of hurdles crossed, as counted manually, were converted to Z-scores and a ratio was used to assess executive function. Animals were tested at the age of 11 and 33 weeks.

#### Puzzle Box

The puzzle box is designed to measure executive function and has been described before in detail^80, 81^. The apparatus has a bright (140 lux), open compartment (60x28 cm) and a dark (∼0 lux), closed compartment (15x28 cm), divided by a partition with an opening (width: 4 cm) in it. The mouse was placed in the dark compartment, and as a result of the natural instinct of mice to avoid open bright areas, would instinctively try to find shelter in the dark compartment by using the opening. To increase the difficulty, the opening can be modified with different materials. Four challenges, with three trials each, were done, divided over 5 days of testing. On each experimental day, the third trial of the current challenge will be done, followed by trials 1 and 2 of the next challenge. Exceptions are the first day, which starts with a habituation trial instead of a third trial, and the fifth day, which only includes the third trial of the fourth challenge. The level of difficulty was increased with each challenge, and challenges were performed in the following order: PVC tunnel (gateway), gateway filled with bedding, tissue plug, and nesting plug. The latency to enter the dark compartment [s] was measured. Animals were tested at the age of 12 weeks.

#### Tripartite Chamber Test

The Tripartite Chamber Test allows for the assessment of social preference^82^. The test apparatus is divided into three chambers (20x40x22 cm each) by transparent Plexiglas partitions, with openings (width: 5 cm) that allow the mouse to visit other chambers. The floor was covered with bedding. During the habituation phase of 5 minutes, the mouse was placed in the central chamber with both entries to the other chambers blocked. During the test phase, wired cages were placed in the outer chambers. One wired cage contained a C3H stimulus mouse of the same age and gender, while the other cage would remain empty. The test mouse was allowed to explore the arena freely for 10 minutes while automated tracking software (Viewer3, Biobserve, Bonn, Germany) recorded the time spent [s], visits to [#], and latency to enter [s] the close proximity of the wire cage. The light intensity in the arena was 130 lux. Animals were tested at the age of 13 weeks.

#### Morris Water Maze

In the Morris Water Maze, spatial learning and memory are assessed. The test was performed as described before^77, 83^. The Morris Water Maze consists of a large circular tank (diameter 120 cm, depth 68 cm), filled with water, in which an escape platform (10x10 cm) is present, submerged 1 cm below the water level. During the first two days (visible task), the mice were trained to find the platform using a visual cue, i.e. a black flag. For the subsequent 8 days (hidden task), mice were trained to find the relocated platform without the visual cue. Extra-maze cues were present to assist the mice in spatial navigation. For the task that took place the last four days (reversal task), the platform was once again relocated and mice were trained to find the platform. Both the hidden task and the reversal task were followed by a day with a probe trial, in which the platform was removed and mice were allowed to swim around freely for 90 seconds. These probe trials were used to assess spatial memory. On every experimental day, except the probe trials, the mouse underwent four trials, each starting from a different location and with the order of entry points changing every day, to ensure dependency on the extra-maze cues. The intertrial interval was 5 minutes, during which the mice were placed in cages on heating platforms to prevent hypothermia. Automated tracking software (Viewer3, Biobserve, Bonn, Germany) measures the latency to reach the platform [s] as well as swimming velocity [cm/s] and total distance traveled [cm]. For the probe trials, time spent in the target quadrant [s], which is where the platform was located during the previous challenge, is recorded. The light intensity in the room was 130 lux. Animals were tested at the age of 19 weeks.

#### IntelliCage

To allow assessment of higher cognitive function in a social setting with minimal experimenter intervention^22^, the IntelliCage was used as described previously^59^. The test environment consists of a type IV cage with the IntelliCage (TSE Systems, Berlin, Germany) placed inside. This IntelliCage is an apparatus with four conditioning corners, which are accessible through a PVC tunnel equipped with a ring radiofrequency identification antenna (RFID), that is capable of identifying individual mice via placed transponders. Each corner contains two water bottles, closed off by motorized doors, that mice can open through a nose poke. The water bottles are equipped with a lickometer, that allows for real-time measurement of the number of licks. During the experiments, mice experience a regular light-dark phase with a maximum light intensity of 80 lux.

The paradigm comprises three challenges: Spatial Learning and Memory, Episodic-Like Memory, and Working Memory (Figure 7b). The three challenges were preceded by a habituation day, during which a nose poke would open every door. Each challenge was followed by an extinction day, during which a nose poke would again open every door, to allow extinction of the previously acquired learning. The first challenge comprised three nights (6pm-6am) during which one corner was accessible to drink from, while the other corners were blocked. During the first night, this was a randomly assigned corner (place learning), with a total of 4 mice assigned per corner. During the second night, this was the diagonally-opposed corner (reversal learning), while during the third night, the allowed corners would switch every three hours between the previously learned corners (multiple reversal learning). The second challenge, which assessed episodic-like memory, consisted of four nights. The acquisition phase happened on the first night, during which the mice were allowed to drink from one assigned corner between 10pm and 12pm, while the other three corners were blocked. During the retrieval phase on the second night, the same corner was allowed in the same time frame. On the third and fourth night (ELM reversal), this pattern would repeat, however, a different corner would be allowed and the time frame would be shifted to 8pm-10pm. The forward shift is to prevent that mice by chance drink in the correct time window if they keep on trying. The last challenge assesses working memory and runs for 4 days during the entire day and night (96 hours). Mice were assigned the corner from which they drank first after activation of the protocol, which would switch clockwise every time they licked in the allowed corner (48 hours). On days 3 and 4 (48 hours), this pattern would switch, as the changing of corners was counter-clockwise. Automated Software recorded the number of visits to each corner, the number of nose pokes, and the number of licks in each corner. Data was analyzed using the Cognition Index of the IntelliR pipeline^59^. This index takes into account the possible errors a mouse could make during a challenge. Animals were tested at the age of 29 weeks.

#### Pheromone-Based Social Box

This pheromone-based social box takes place in the IntelliCage setup and was performed as described before^81^, with minor adaptations. One week after the IntelliCage behavioral phenotyping, mice were placed again in the IntelliCage set-up (light intensity: 80 lux). The home cage set-up was expanded with on each side a smaller box connected via Plexiglas tubes (4 cm diameter). The Plexiglas tubes were equipped with RFID antennas. The experimental set-up comprised three phases. During the pre-habituation phase, animals were allowed to re-habituate to the IntelliCage for 1.5 hours. Subsequently, the habituation phase started, in which mice were allowed to freely explore both the home cage and the two social boxes for one hour. Both social boxes were filled with fresh bedding to prevent side preferences. The test phase followed a similar set-up as the habituation phase, with the difference that during this phase, one of the boxes was filled with fresh bedding while the other box was filled with used bedding from C3H mice from the opposite gender. Time spent in each social box [s] was used for further analysis. Animals were tested at the age of 32 weeks.

#### Pre-Pulse Inhibition

To assess sensorimotor gating, Pre-Pulse Inhibition was executed, as described before^77, 83^. In short, mice were placed inside a small metal cage (85x40x40 mm) to restrict movement. This cage was placed on a movement-sensitive platform in a dark (∼0 lux), sound-attenuating chamber. The experimental sessions lasted for 20 minutes, during which acoustic stimuli were given to provoke a startle response. First, habituation to 65 dB white noise was provided for the mice, after which acoustic stimuli were provided. Either a startle pulse (120dB) was provided alone, or it was preceded by a pre-pulse (70, 75, 80dB) which would then be given 100ms before the startle pulse. Each acoustic stimulus or combination thereof was presented 10 times. The startle response was averaged for each intensity per mouse and the pre-pulse inhibition (PPI) was calculated as a percentage of the startle response, using the following formula:

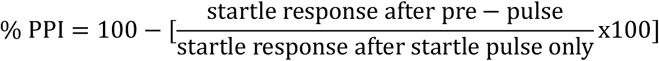

Animals were tested at the age of 33 weeks.

#### Detailed acoustic assessment

The auditory system function was assessed through the analysis of a detailed tone-intensity-startle response curve^82^. Using a similar set-up as the Pre-Pulse Inhibition test, mice were submitted to auditory stimuli. For a detailed assessment of auditory function, the acoustic stimuli ranged from 65 dB to 120 dB (in steps of 3 dB), given in a pseudo-randomized order. The experimental session lasted 50 minutes, during which each intensity was presented 10 times. The startle response was averaged for each intensity per mouse. Animals were tested at the age of 33 weeks.

#### Mental Time Travel

To assess higher cognitive function in mice, Mental Time Travel was developed, as described in detail by^84^. In short, the Mental Time Travel test takes place in the IntelliCage setup and lasts nine days. In comparison to the IntelliCage behavioral phenotyping as described above, the Mental Time Travel test uses an air puff of 1.5 bar as a type of negative reinforcement to punish drinking attempts at incorrect corners. During the test days, always one corner will be deemed incorrect while the other three corners allow access to water. These corners are randomly assigned, with a maximum number of 4 mice per corner. The test started with a place learning day, similar to the place learning day in the IntelliCage behavioral phenotyping. This is followed by a training phase of four days, in which access to water was limited in one corner from 6pm-8pm and an attempt to drink from the incorrect corner was punished with an air puff. The incorrect, punished corners follow a distinct pattern: the first assigned punished corner (day two), the diametrically opposed corner of day two (day three), the horizontally opposed corner of day three (day four), and the diametrically opposed corner of day four (day five). After this, the test phase took place from day 6 to day 9, following the same pattern. For the analysis, the corners were assigned a label, based on the recentness of the punishment: “punished” (n=0 days after punishment), “recent” (n=1 days after punishment), “intermediate” (n=2 days after punishment), and “safe (n=3 days after punishment). The averaged percentages of visits to each corner were used for further statistical analysis. Animals were tested at the age of 34 weeks.

#### Fear Conditioning

The Fear Conditioning test has been performed as before, with minor adaptations^77^. The Fear Conditioning comprised three phases: training, context and cued fear conditioning. The experiment took place within a computerized fear conditioning system. The system includes a sound-insulated cabinet (73x64x36 cm), which contained a conditioning chamber with a Plexiglas door (light intensity 15 lux). For the training and context phases, a rectangular chamber (36x20x20 cm) with a shock grid floor (composed of stainless-steel bars) was used with a light intensity of 114 lux. The cued fear conditioning phase took place in a novel, half-circular chamber (25x20x15 cm), with a Plexiglas door and white floor, and a light intensity of 11 lux. The training phase consisted of a habituation phase in which no stimuli were given, to establish a baseline for freezing behavior. Afterward, an acoustic stimulus (conditioned stimulus, CS, 75 dB) was given, immediately followed by an electric foot shock (unconditioned stimulus, US, 0.4 mA, 2s). This pattern was repeated once, followed by a period of 30 seconds in which no stimulus was given, to avoid mice making the association of the aversive event to handling by the experimenter. The context session took place 72h later, and comprised mice being exposed to the same context; however, no stimuli were provided. Four hours later, the cued session took place, in which mice were placed in the novel environment and exposed to a protocol starting with a 120-second stimulus-free (pre-cue) period immediately followed by a 120-second period in which the mice were exposed to the CS (cue period). During all three sessions, the duration of freezing [s], in which freezing is defined as a complete lack of movement, was automatically recorded using a video camera and software (Video Freeze, MED associates, St. Albans, Vermont, USA). Animals were tested at the age of 37 weeks.

#### Complex Running Wheel

To test motor performance and learning, a complex running wheel set-up was used as described before^85^. In short, mice were individually housed in type III cages for 21 consecutive days. These cages contained a complex running wheel, where random bars were omitted (16 out of 35). Running was voluntary, and mice had *ad libitum* access to food and water. Software was used to monitor running behavior per night, measuring the following parameters: total distance [s], time spent on the wheels [s], maximal velocity, and average velocity [s]. During the course of the test, mice experienced a regular light-dark phase with intensities between ∼0 and 190 lux. Animals were tested at the age of 38 weeks.

#### Magnetic Resonance Imaging (MRI)

After perfusion at the age of 55 weeks, brains from male *CnpCre::EPOR* mice were extracted and fixed with 4% formaldehyde, 2.5% glutaraldehyde and 0.5% NaCl in phosphate buffer (pH 7.4)^86^. After 1 week, brains were transferred to 1% PFA in 0.1M phosphate buffer for MRI. Post-mortem MRI was conducted at a magnetic field strength of 9.4T using a 4-channel receive-only mouse head coil (Biospec, Bruker BioSpin, Ettlingen, Germany). For volumetric analyses, magnetization transfer (MT)-weighted images were acquired with a 3D fast low-angle shot (FLASH) sequence (TE/TR = 3.4/15.2ms, flip angle 5°, Gaussian-shaped off-resonance pulse (off-resonance frequency 7.5ppm, RF power 6µT), and an isotropic spatial resolution of 100µm. The MT-weighted images were converted to the NIfTI format, denoised and bias field corrected^87^. The signal of the surrounding fluid was manually removed, an unbiased anatomical study template generated and voxel-wise Jacobian determinants were calculated from nonlinear deformation fields (twolevel_ants_dbm, https://github.com/CoBrALab/). Student’s t-test was performed on the 3D Gaussian-smoothed maps of Jacobian determinants using the 3dttest++ function of AFNI. To visualize volume differences between mutants and controls, z-scores smaller than -2.57 and larger than 2.57 were overlaid on the study template. To quantify the volume of specific brain areas regions of interest (ROIs), including corpus callosum, hippocampus, fimbria, cerebellum, olfactory bulb, striatum, and cortex (without hippocampus) were determined on the study template by manual segmentation using ITK-SNAP (Version 3.8.0). ROIs were subsequently retransformed into the subject space, individually checked and manually corrected if necessary. The final volume information was then extracted for statistical analysis.

#### Electron Microscopy (EM) Analyses

After performing the MRI scan, the fixed brains from male *CnpCre::EPOR* mice were used for EM. Sagittal sections (300µm) were cut with vibratomes. Fimbria, corpus callosum and hippocampus were separated and embedded in epoxy resin (Serva, Heidelberg, Germany) as described^88^. Ultrathin (60 nm) sections were prepared on a PTPC Powertome Ultramicrotome (RMC, Tucson, AZ, USA) with a diamond knife (Diatome AG, Biel, Switzerland). Ultrathin sections were attached on forward-coated copper grids (AGAR Scientific, Essex, UK) and stained with UranyLess (Electron Microscopy Science, Hatfield, Panama) for 20min to improve the contrast. Sections were washed with ddH_2_O 3x before imaging with Zeiss EM900. Non-overlapping 10-15 random images of fimbria, corpus callosum and CA1 region of hippocampus were taken per animal at 4000x. All image analyses were performed on Fiji (ImageJ 1.53 t). Myelinated axons were counted by cell counter function for CA1 region (500 µm^2^), corpus callosum (200 µm^2^) and fimbria (200 µm^2^), 5 images per sample. To quantify axon diameter of corpus callosum and fimbria, 25-30 axons were chosen randomly from each image (5 images per sample) and the area of each axon was measured by drawing the ROI. The g-ratio was calculated as ratio between axonal diameter and the outer diameter of the corresponding myelin sheath.

#### Single-Nuclei RNA Sequencing

For the current dataset, 23 male mice on P49 (*N* EPO = 11, *N* placebo = 12), 24h after the last rhEPO/placebo injection, were sacrificed by cervical dislocation. The brain was immediately removed, with the right hippocampus dissected on an ice-cold plate, immersed in liquid nitrogen, and stored at -80°C. Two right hippocampi of the same treatment group were collected in one tube for sequencing (one tube in the rhEPO group with only one right hippocampus). Final analysis was performed on *N*=6 tubes *per* group. Single nuclei suspension was prepared using 10x Genomics Chromium Single Cell 3’ Reagent Kits v3 (10X Genomics, Pleasanton, CA) according to manufacturer’s protocol. Quantity and quality of cDNA were assessed by Agilent 2100 expert High Sensitivity DNA Assay. cDNA samples were sequenced on NovaSeq 6000 S2 flowcell at UCLA Technology Center for Genomics and Bioinformatics. Raw and processed single-nuclei RNA sequencing (snRNA-seq) data are publicly available on GEO via accession code GSE220522.

#### Single-Nuclei RNA-Seq Data Processing

The complete snRNA-seq dataset used in this manuscript was first processed and described in another study of our group^8^. Briefly, FASTQ files were mapped to the mouse genome (version mm10) using CellRanger (v6.1.1; 10x Genomics) to obtain gene/nucleus count matrices. The corresponding genome reference and gene transfer format (GTF) files were sourced from Ensembl and processed using the mkref function of CellRanger. The alignment procedure was executed using the standard parameters as outlined in the developer’s manual. Subsequently, to address possible issues like batch effects and differential gene expression, background RNA was eliminated using CellBender version 0.2.1106. The quality of the alignment and data matrices were evaluated using the downstream processing utilities provided by CellRanger. The processed data set from our previous work^8^ was used as the starting point for the analysis described here. The first step was to subset the major cluster identified as “Oligodendrocytes” using Seurat (v4.3.0)^43^, implemented in R (v4.3.0)^89^. Then, the resulting dataset went through the following steps: (1) normalization using the standard Seurat procedure, (2) the ‘find variable features’ process with standard settings, (3) data scaling regressing mitochondrial and ribosomal percentages, and cell cycle scores, (4) and Principal Component Analysis using Seurat’s standard settings. Afterwards, we ran Harmony (v 1.0.1)^90^ in the resulting object, followed by nearest neighbors calculation and cluster identification using Seurat’s shared nearest neighbor (SNN) modularity optimization based clustering algorithm^43^. UMAP dimensionality reduction was applied and used for data visualization. To ensure the clusters sent for downstream analysis contained only nuclei of interest, we checked them for expression of key OL lineage markers (*Apc*, *Bcas1*, *Bmp4*, *Cnp, Cpsg4, Mog, Olig1*, *Olig2*, *Pdgfra*, *Plp1*, *Sox6*, *Sox10*). These steps were repeated until all clusters that did not express OL lineage markers were removed. Monocle3 (version 1.3.1)^42^ was then used to calculate a pseudotime measurement using principal graph algorithm together with the previously calculated UMAP. This pseudotime measurement represents the current state of each nucleus in the transcriptional trajectory of the dataset. Combinations of the established OL lineage markers and pseudotime were used to determine the final identity of the clusters.

Differential gene expression between EPO and placebo samples was calculated using the FindMarkers function of Seurat^43^ using its standard settings (±0.25 average log2 fold change, adjusted p-value<0.05) with the likelihood-ratio test for single cell gene expression (“bimod”)^91^. Differentially expressed genes (DEG) found for each cluster were used in WebGestalt^44^ to calculate Gene Ontology (GO) terms for biological processes (“no redundant”) using default settings in the Over Representation Analysis. The significance level was set to FDR<0.05 calculated through the Benjamini-Hochberg (BH) procedure and the background to ‘genome’.

Finally, we employed SCENIC^56^ to analyze gene regulatory networks (GRN) using the same basic steps described in our prior work^8^. Briefly, GENIE3^92^ was used to identify co-expression networks via genome-wide correlation analysis. This was followed by the inference of regulatory networks from expression data using random forest regressions to construct the modules, which are sets of co-expressed genes with a master regulator. Only modules with a significant threshold of predicted weight for each module regulator (links with weight >0.005) were used. These modules were further filtered to keep only those with a minimum of 20 genes for the AUCell scores (with aucMaxRank=10%). AUCell quantified AUC values were subsequently transformed into a binary activity matrix as recommended by SCENIC^56^. The genes that constructed regulons exclusive to either EPO or placebo treatments, minus lncRNAs or pseudogenes, were used to calculate GO enrichment using clusterProfiler (version 4.10.0)^57^ standard settings and at least two genes per enriched GO term. Detailed codes for all analyses and plots for this section are available on GitHub under the following link: https://github.com/vgastaldi/Ye_Gastaldi_et_al_EPO_snRNAseq_Oligo or at the following Zenodo repository: https://doi.org/10.5281/zenodo.1580666493.

#### Statistical Analyses

Statistical analyses were performed using Prism10 software (GraphPad Software) and R (The Comprehensive R Archive Network). All values represent mean±SEM (standard error of the mean) with N numbers (i.e. number of mice/group) and statistical tests specified in the text or corresponding figure legends. Normal distribution of data was assessed using the Shapiro-Wilk test. For comparison of two groups, student’s unpaired two-tailed t-tests were performed in normally distributed data and two-tailed Mann-Whitney U test in non-parametric data. Multiple groups were compared using one-way ANOVA for normally distributed data or Friedman’s test for non-parametric data for analysis of immunohistochemical data. For analysis of behavioral data, the multiple group comparisons were done by performing a repeated measure two-way ANOVA for normally distributed data or the two-way Friedman’s test for non-parametric data; p-values <0.05 were considered statistically significant.

## Data Availability

The single-nuclei RNA sequencing (snRNA-seq) data are publicly available on GEO via accession code <u>GSE220522</u>. Source data are provided with this paper.

## Code Availability

All code used for the bioinformatics analysis of this work is available at the following Zenodo repository: https://doi.org/10.5281/zenodo.1580666493 or alternatively at https://github.com/vgastaldi/Ye_Gastaldi_et_al_EPO_snRNAseq_Oligo.

## Acknowledgements

This work has been supported by the European Research Council (ERC) Advanced Grant to HE under the European Union’s Horizon Europe research and innovation programme (acronym *BREPOCI;* grant agreement No 101054369), as well as by the Max Planck Society, the Max Planck Förderstiftung, the Deutsche Forschungsgemeinschaft (DFG, German Research Foundation), via DFG-Center for Nanoscale Microscopy & Molecular Physiology of the Brain (CNMPB). Research in the labs of HE and KAN is funded by DFG TRR-274 /1 2020 – 408885537. LY has obtained funding from the National Natural Science Foundation of China (82100091). VDG received support from the IMPRS-Genome Science PhD program. YC is recipient of a grant from the Peter and Traudl Engelhorn Foundation. XY was supported by the China Scholarship Council (202106160020). Research in the labs of DG, RK and KAN is supported by the Adelson Medical Research Foundation.

## Author Contributions

Concept, design, supervision: HE,KAN

Funding acquisition: HE,RK,DG,KAN

Drafting manuscript and display items: LY,VDG,YC,AFW,XY,HE

Data acquisition/generation: LY,VDG,YC,AFW,XY,SE,MH,AR,QW,UJB,WM,SB

Data analyses & interpretation: LY,VDG,YC,AFW,XY,HE,KAN,RK,DG,MS,WM,SB

**All authors read and approved the final version of the manuscript.**

## Competing Interests

The authors declare no competing financial or other interests in connection with this article.

---

**Supplementary Table 1:** Complete list of differentially expressed genes. List of all differentially expressed genes (DEG) obtained using the ‘bimod’ test in Seurat with the following settings: minimum average log2FC ±0.25; adjusted p-value≤0.05, ‘bimod’ test, multiple correction done using Bonferroni. The colors of the clusters match the ones used in Figure 3a.

**Supplementary Table 2:** Enriched Gene Ontology terms using differentially expressed genes. Complete list of enriched Gene Ontology (GO) terms for biological processes obtained using the DEG calculated through Seurat’s ‘bimod’ test. Standard settings for Over Representation Analysis were used in WebGestalt, with FDR<0.05 (Benjamin-Hochberg) and background set to genome.

**Supplementary Table 3:** Complete list of regulon modules calculated by SCENIC. Regulons obtained after SCENIC analysis. Only links with weight > 0.005 were kept, following filtering to keep modules with a minimum of 20 genes prior AUCell score calculation. The colors of the clusters match the ones used in Figure 3a.

**Supplementary Table 4:** Enriched Gene Ontology terms calculated using SCENIC regulons. Complete list of enriched Gene Ontology (GO) terms calculated with clusterProfile 4.0 using the regulon modules obtained through SCENIC. These were calculated by over-representation analysis (adjusted p-value <0.05, Benjamini-Hochberg procedure) and are divided between regulons exclusively active in rhEPO or Placebo treatments. The colors of the clusters match the ones used in Figure 3a.

**Supplementary Table 5:** Detailed overview of the behavioral assessment of mice with EPOR deletion in MOL. Mice with EPOR deletion in MOL underwent an extensive behavioral battery between the ages of 9 weeks and 50 weeks. Data was analyzed using two sample t-test with Welch’s correction (t) or Mann-Whitney U-test (U), dependent on the data distribution. Two-way ANOVA or two-way Friedman, dependent on the data distribution, was done for comparison across time or test phase.

## Notes

### Competing Interest Statement

The authors have declared no competing interest.

